# DNA-templated spatially controlled proteolysis targeting chimeras for CyclinD1-CDK4/6 complex protein degradation

**DOI:** 10.1101/2024.09.18.613743

**Authors:** Rong Zheng, Abhay Prasad, Deeksha Satyabola, Yang Xu, Hao Yan

## Abstract

Constraining proximity-based drugs, such as proteolysis-targeting chimeras (PROTACs), into its bioactive conformation can significantly impact their selectivity and potency. However, traditional methods for achieving this often involve complex and time-consuming synthetic procedures. Here, we introduced an alternative approach by demonstrating DNA-templated spatially controlled PROTACs (DTACs), which leverage the programmability of nucleic-acid based self-assembly for efficient synthesis, providing precise control over inhibitors’ spacing and orientation. The resulting constructs revealed distance- and orientation-dependent selectivity and degradation potency for the CyclinD1-CDK4/6 protein complex in cancer cells. Notably, an optimal construct DTAC-V1 demonstrated the unprecedented synchronous degradation of entire CyclinD1-CDK4/6 complex. This resulted in the effective cell cycle arrest in G1 phase, and further therapeutic studies showed its potent anti-tumor effects compared to inhibitors alone. These findings present a novel framework for PROTACs design, offering critical insights that may inform the development of other proximity-induced therapeutic modalities.

## Introduction

Controlling biomolecular proximity with functional materials offers great potential for regulating intracellular processes and the advancement of new therapeutic approaches. In this context, PROteolysis TArgeting Chimeras (PROTACs), a class of proximity induced protein degraders, hold significant transformative potential by selectively targeting disease related proteins^1,2^. Since the invention of the first PROTAC in 2001^3^, critical breakthroughs have been achieved, including the degradation of previously undruggable targets like STAT3^4^, and various transcription factor proteins^5^. Despite initial success, significant hurdles remain in the design of various PROTACs, which require optimization of linker length, time-demanding and labor-intensive synthesis for each candidate^6^. Moreover, constraining PROTACs in their bioactive conformation remains challenging and underexplored^7,8^. Programmable nucleic acid-based scaffolds provide a promising solution towards the design and optimization of advanced constructs due to their programmability, biocompatibility and high-throughput potential^9^. Previous studies have demonstrated that nucleic acid-based modalities could be utilized as substrates for PROTAC, such as RNA-PROTAC^10^, TF-PROTACs^11,12^, aptamer-PROTAC^13,14^, Z-DNA-PROTAC^15^, and DbTAC^16^. Nonetheless, these systems were unable to precisely control both the ligand spacing and orientation, which may impact protein degradation rate and selectivity, and are often limited to proteins that recognize specific oligonucleotide sequences and motifs.

In this work, we introduced **<u>D</u>**NA-templated spatially controlled proteolysis-**<u>TA</u>**rgeting **<u>C</u>**himeras (**DTACs**) by integrating small molecule inhibitors of the cullin-RING E3 ubiquitin ligase (CRL4^CRBN^) and the protein of interest (POI) with a customizable DNA duplex scaffold. In contrast to traditional PROTACs design, this study highlights several advantages: 1) The spacing and orientation between inhibitors can be adjusted using the programmable DNA scaffold without altering the overall chemical composition, thus reducing complex synthetic routes; 2) Multiple combinations of distance and orientation libraries of DTACs can be generated by reusing a specific ssDNA-inhibitor across different versions, obviating the need for synthesis each time a new version is developed; 3) Rapid self-assembly of different DTAC versions can be achieved at high concentration with high fidelity of self-assembly. Overall, the ease of oligonucleotide synthesis and functionalization, combined with the simplicity and scalability of DTAC self-assembly, facilitates the rapid design of chimeric molecules, leading to faster development and optimization.

To evaluate the DTAC constructs, we targeted Cyclin-dependent kinases 4 and 6 (CDK4/6) and their activating partners, D-type cyclins, which play a pivotal role in cell cycle regulation^17^. The Cyclin D-CDK4/6 complex drives tumorigenesis in several cancers by promoting uncontrolled cell proliferation^18,19^. Inhibiting CDK4/6 has shown clinical success, particularly in treating hormone receptor-positive breast cancers^20,21^, but resistance to CDK4/6 inhibitors remains a significant issue^22,23^. Emerging research suggests that targeting Cyclin D1-CDK4/6 complex degradation, rather than solely inhibiting CDK4/6, may offer a broader therapeutic impact by eliminating both CDK4/6-dependent and independent roles in tumor progression^24^. However, targeting Cyclin D1 (encoded by CCND1) remains challenging due to the absence of a catalytic pocket or active site^25^. Several small molecule-based PROTACs, including CP-10^26^, MS140^27^, and BSJ-03-123(BSJ)^28,29^ have been made to target this complex. Nonetheless, these approaches reported only selective degradation of CDK4 or CDK6, or both CDK4 and CDK6, while leaving Cyclin D1 unaffected. Recently, a study employing a bridge PROTAC method demonstrated preferential degradation of Cyclin D1^30^, yet, to our knowledge, no method has successfully achieved synchronous degradation of the entire CyclinD1-CDK4/6 protein complex.

Here, we found that with optimal spacing and orientation of inhibitors, a construct (DTAC-V1) can induce synchronized degradation of the entire Cyclin D1-CDK4/6 complex. Furthermore, we demonstrated that the spatial arrangement of the inhibitors is critical in modulating both the selectivity and degradation efficacy. Proteomic and anti-proliferation studies further highlighted the downstream effects of Cyclin D1-CDK4/6 degradation, revealing a potent anti-tumor growth response. These findings underscore the potential of DTAC to address key limitations associated with conventional PROTACs strategies.

## Results and discussion

### Synthesis of the DTAC distance and orientation library

The fundamental DTAC design consists of two primary components: a customizable DNA duplex scaffold and two small molecule inhibitors that target the E3 ligase (termed as E3i) and the POI (termed as POIi). To integrate these components, we employed solid-phase oligonucleotide synthesis, chemical synthesis of small molecules, and strategic oligo modification and conjugation techniques. The DNA duplex was chosen for the DTAC formulation due to its rigid and predictable helical structure with a corresponding persistent length of approximately 50 nm that could provide an ideal scaffold with the ability to tune the distance and orientation of inhibitors^31^. As an initial test module, we utilized a 20 base pairs DNA duplex that offered a tunable distance of ∼1-6 nm between the inhibitors. In the DTAC duplex, strand A is designated for conjugation to the E3i and strand B is designated for conjugation to the POIi, at selective sites. First, an amino-modifier serinol phosphoramidite was incorporated at a specific internal site of both constituent strands through solid-phase oligo synthesis. This site was subsequently used to conjugate a dibenzocyclooctyne (DBCO) group through an amine reactive NHS-ester reaction (**Figure S7A**), which further enabled the attachment of inhibitors via strain-promoted alkyne-azide cycloaddition (SPAAC) click chemistry, as illustrated in **Figure 1A**. To enhance the stability of DTAC in vitro, both strands A and B were modified with three terminal phosphorothioates during oligo synthesis.

**Figure 1:**
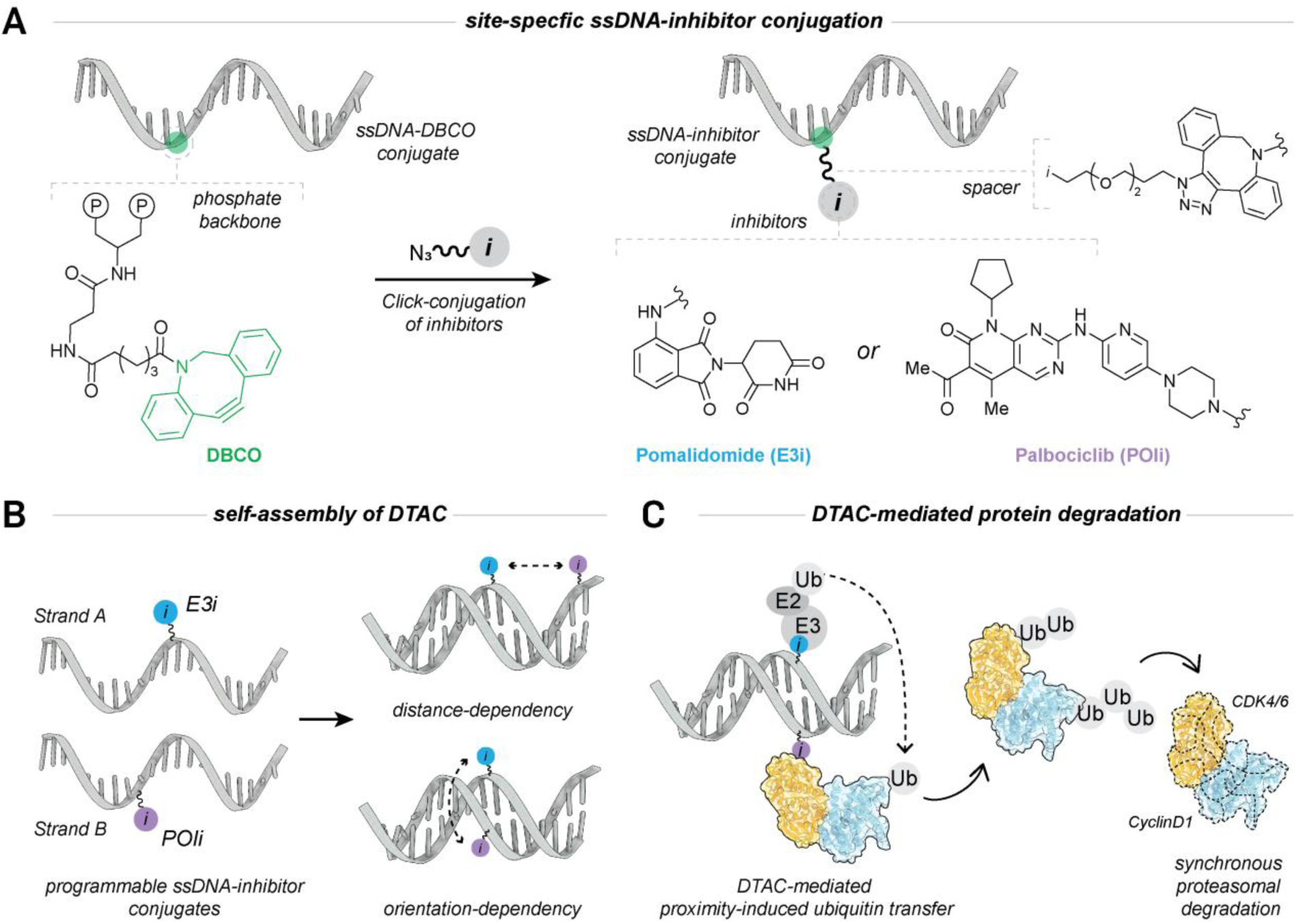
Overview of DTAC synthesis and protein complex degradation. **(A)** The site specifically modified ssDNA with DBCO group was reacted with azido-PEG2-small molecule inhibitors through SPAAC click chemistry. (**B)** The DTAC platform was constructed by thermally annealing two complementary ssDNA-inhibitor (strand A with E3i and strand B with POIi) conjugates. The distance or orientation DTAC library was generated by mixing different combination of strand A and B. (**C)** The optimal conformation and distance between inhibitors on DNA scaffold induces selective and efficient degradation of CyclinD1-CDK4/6 complex.

We selected two small molecule inhibitors for the current study: pomalidomide (POM) an E3i targeting the CRL4^CRBN^ ubiquitin ligase complex (REFs), and palbociclib (PB), which serves as a POIi that targets both CDK4, and CDK6. The inhibitors were modified with a hydrophilic PEG-2-azide handle (**Figure S1, S4**), and reacted them with the DNA-DBCO conjugates in a 1:2 ratio, whereby successful conjugation was observed with a yield of approximately 95%. The DNA-inhibitor conjugates were purified via HPLC and characterized by quadrupole/time-of-flight mass spectrometer (Q-TOF MS) to confirm the expected molecular weight of the desired products (**Figure S8, S9**).

Lastly, the modified ssDNA-inhibitor conjugates (Strand A with E3i and Strand B with POIi) were self-assembled in a 1:1 ratio in 1xPhosphate buffered saline (PBS, pH 7.4) using a 3min in rapid annealing from 80°C to 25°C to form a DTAC library with controlled distances and orientations **(Figure 1B)**. We designed four distance versions namely DTAC-V1, DTAC-V2, DTAC-V3 and DTAC-V4 to systematically investigate the effects of inhibitors’ spacing, with distances ranging from ∼1 nm to 6 nm. Additionally, five different constructs (DTAC-V5 to DTAC-V9) were designed to evaluate the orientational effects by varying the position of E3i on strand A. Notably a common strand B with POIi inhibitor was utilized across three distance versions and all orientational versions of DTAC library (**Table S2**). Similarly, a common strand A with E3i inhibitor can be opted for while screening different E3 ligases against the protein target. Successful self-assembly of the DTAC library was assessed by an electrophoretic mobility shift assay (EMSA) using native-PAGE, which showed higher retention of the DTAC library compared to DNA duplex only control (**Figure S10**). All constructs were rapidly assembled with high fidelity of self-assembly, even at concentrations up to 50 µM.

### Distance and orientation-dependent degradation of CyclinD1-CDK4/6

Having established the DTAC platform, we next evaluated its distance- and orientation-dependent selectivity and degradation rates in U251 cancer cells using Western blot (WB) analysis. As a proof of concept, we targeted the Cyclin D1-CDK4/6 complex (CyclinD1-CDK4 and CyclinD1-CDK6). Due to the affinity of the Palbociclib (PB) for CDK4 and CDK6, we were initially interested in whether the different DTAC variants could differentiate between these two proteins. To assess this, cells were transfected with four DTAC variants (DTAC-V1, DTAC-V2, DTAC-V3, and DTAC-V4 as shown in **Figure 2A** at concentrations of 20 and 100 nM. Remarkably, DTAC not only degraded CDK4/6 but also the entire CyclinD1-CDK4/6 complex, with each variant exhibiting distinct degradation selectivity (**Figure 2B**). DTAC-V1 showed excellent degradation of the entire CDK4/6-Cyclin D1 complex, achieving degradation rates at 100 nM of ∼60% for CDK6, and ∼90% for both CDK4 and Cyclin D1, respectively. By contrast, DTAC-V2 preferentially degraded the CDK4-Cyclin D1 complex, with degradation rates up to 70% for CDK4 and 50% for Cyclin D1, with only a minimal (∼20%) effect on CDK6, thus exhibiting preferential selectivity for the CDK4-Cyclin D1 complex. Interestingly, DTAC-V3, with a 2 nm inhibitors’ spacing, preferentially targeted CDK4, while having minimal impact on CDK6 and Cyclin D1(**Figure 2C**). This selectivity possibly attributed to optimal spacing between inhibitors that influence CDK4 facing with E3 ligase and facilitate its preferential polyubiquitination. Furthermore, DTAC-V4, with inhibitors’ spacing of ∼6 nm exhibited no significant impact on the protein complexes. While screening different DTAC versions, we utilized a DNA-only scaffold (**Table S1**) as a control, which exhibited no impact on protein complex expression level even at 100 nM. Further controls confirmed by both WB and immunofluorescence (IF) staining (**Figure S12**) using the DNA scaffold with either E3i (DNA-E3i) or POIi (DNA-POIi) alone yielded no effect on protein degradation. Altogether, our findings demonstrate that tuning the spacing between inhibitors on the DNA duplex allows distance-dependent selectivity and degradation rates. However, even though DTAC-V1 and DTAC-V2 have similar ligand spacing (∼1.2 nm), they exhibit different selectivity and degradation potency. The optimal PROTAC conformation can minimize the entropic penalty for achieving a bioactive state between the E3 ligase and the protein target ^8^. Therefore, this led us to investigate whether the different conformation of inhibitors in DTAC-V1 and DTAC-V2 affect the degradation rate and selectivity of the protein complex. To evaluate this, we transfected cells with five different versions of the orientational library (DTAC-V5 to DTAC-V9) as shown in **Figure 2D**. WB analysis revealed a gradual increase in CDK6 degradation rates at 100 nM as the conformation changed from DTAC-V5 to DTAC-V9 (**Figure 2E, 2F**). However, no significant differences in activity were observed for the CDK4-CyclinD1 complex across DTAC-V5 to DTAC-V8. As anticipated from the previous distance library analysis, DTAC-V9 (same construct as DTAC-V1) demonstrated consistent activity at both 20 and 100 nM. These results suggest that changing orientation from DTAC-V5 to DTAC-V9 enhances selectivity for CDK6 while maintaining efficient degradation of CDK4-CyclinD1 across all versions.

**Figure 2.**
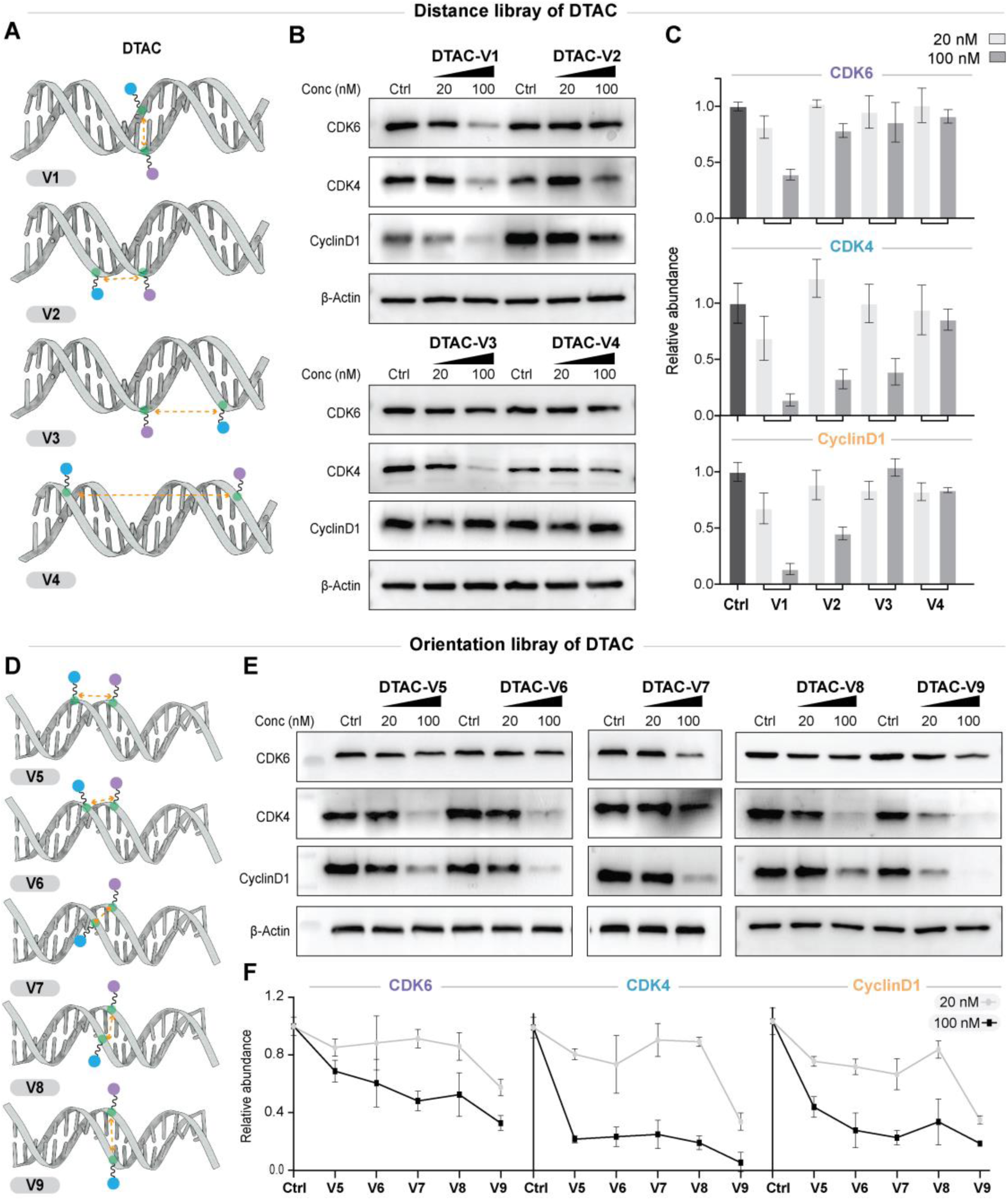
Distance- and orientation-dependent selectivity and degradation of the Cyclin D1-CDK4/6 complex. **(A)** Schematic representation of the DTAC distance library. (B) Western blot (WB) analysis showing Cyclin D1 and CDK4/6 degradation in U-251 cells treated with the indicated DTAC variants at the specified concentrations for 14 hours. Results are representative of three independent experiments. (C) Quantification of relative Cyclin D1, CDK4, and CDK6 levels compared to the control group following treatment with the distance-dependent DTACs. Data shown are the mean values ± SD from three independent experiments. (D) Schematic representation of the DTAC orientation library. (E) WB analysis showing Cyclin D1 and CDK4/6 degradation in U-251 cells treated with the indicated orientation variants (DTAC-V5 to DTAC-V9) at the specified concentrations for 14 hours. Results are representative of three independent experiments. (F) Quantification of relative Cyclin D1, CDK4, and CDK6 levels compared to the control group following treatment with the orientation-dependent DTACs. Data shown are the mean values ± SD from three independent experiments.

### DTAC-V1 mediated synchronous degradation of CyclinD1-CDK4/6

To determine whether the observed degradation of CyclinD1 is a consequence of CDK4/6 degradation or if all three proteins are degraded simultaneously, we conducted a time-course WB analysis for all three proteins following treatment with DTAC-V1 (**Figure 3A**). Notably, Cyclin D1, CDK4, and CDK6 protein levels began to decrease as early as 4 h post-treatment, with significant reductions becoming evident by 6 h **(Figure 3B)**. These reductions persisted, with all three proteins exhibiting further degradation at low levels beyond 12 h, which remained consistent when tested up to 48 h **(Figure 3B).** Additionally, DTAC-V1 showed a concentration-dependent degradation of the CyclinD1-CDK4/6 complex in U-251 cells, with CDK6 exhibiting a DC_50_ of 55 ± 18 nM and a D_max_ of 70 ± 6%, CDK4 showing a DC_50_ of 12 ± 5 nM and a D_max_ of 94 ± 3%, and Cyclin D1 displaying a DC_50_ of 20 ± 13 nM with a D_max_ of 91 ± 1% at 16 h **(Figure 3C, D)**. Given that Cyclin D1 is also regulated by the downstream Rb-E2F signaling axis, we next investigated the impact of DTAC-V1 on CCND1 and CDK4/6 mRNA expression levels using quantitative reverse transcriptase polymerase chain reaction (RT-qPCR) as shown in **Figure 4A**. Neither DTAC-V1 nor the control compounds DNA-POIi or DNA-E3i indicated significant changes in CCND1, CDK4, or CDK6 mRNA levels, suggesting that the observed depletion of Cyclin D1 occurs via post-transcriptional mechanisms. These findings collectively demonstrate that DTAC-V1 induces a synchronous degradation of the entire Cyclin D1-CDK4/6 complex, distinct from a purely selective degradation of CDK4/6 alone.

**Figure 3.**
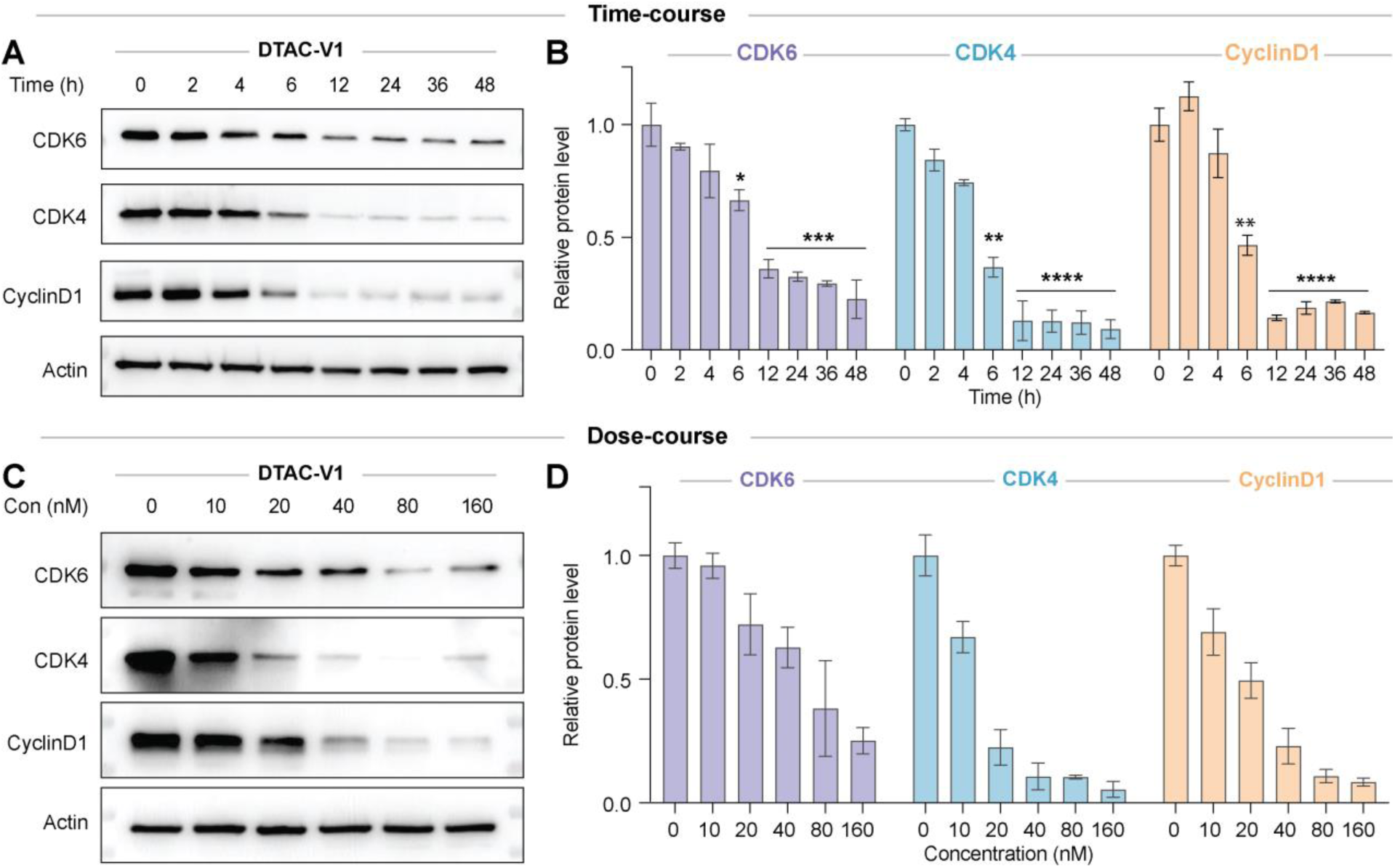
Dose- and time-course degradation of CyclinD1-CDK4/6. **(A)** Time-course analysis of CDK6, CDK4, and Cyclin D1 protein degradation induced by DTAC-V1. U-251 cells were treated with DTAC-V1 at the indicated time points, and protein levels were assessed. Data are representative of three independent experiments. **(B)** Quantification of relative Cyclin D1, CDK4, and CDK6 protein abundance at each time point following treatment with 100 nM DTAC-V1. Data are presented as the mean ± SD from three independent experiments. P-values were calculated relative to protein abundance at 0 h. Statistical significance is indicated as ****P < 0.0001, ***P < 0.001, **P < 0.01, and *P < 0.05. The horizontal lines above the bars indicate that the grouped bars share the same P-value. **(C)** DTAC-V1 induces a concentration-dependent degradation of the Cyclin D1-CDK4/6 complex. U-251 cells were treated with DTAC-V1 at the indicated concentrations for 12 hours. Results are representative of two independent experiments. **(D)** Quantification of Cyclin D1, CDK4, and CDK6 protein levels relative to the control group following DTAC-V1 treatment from two independent experiments.

**Figure 4.**
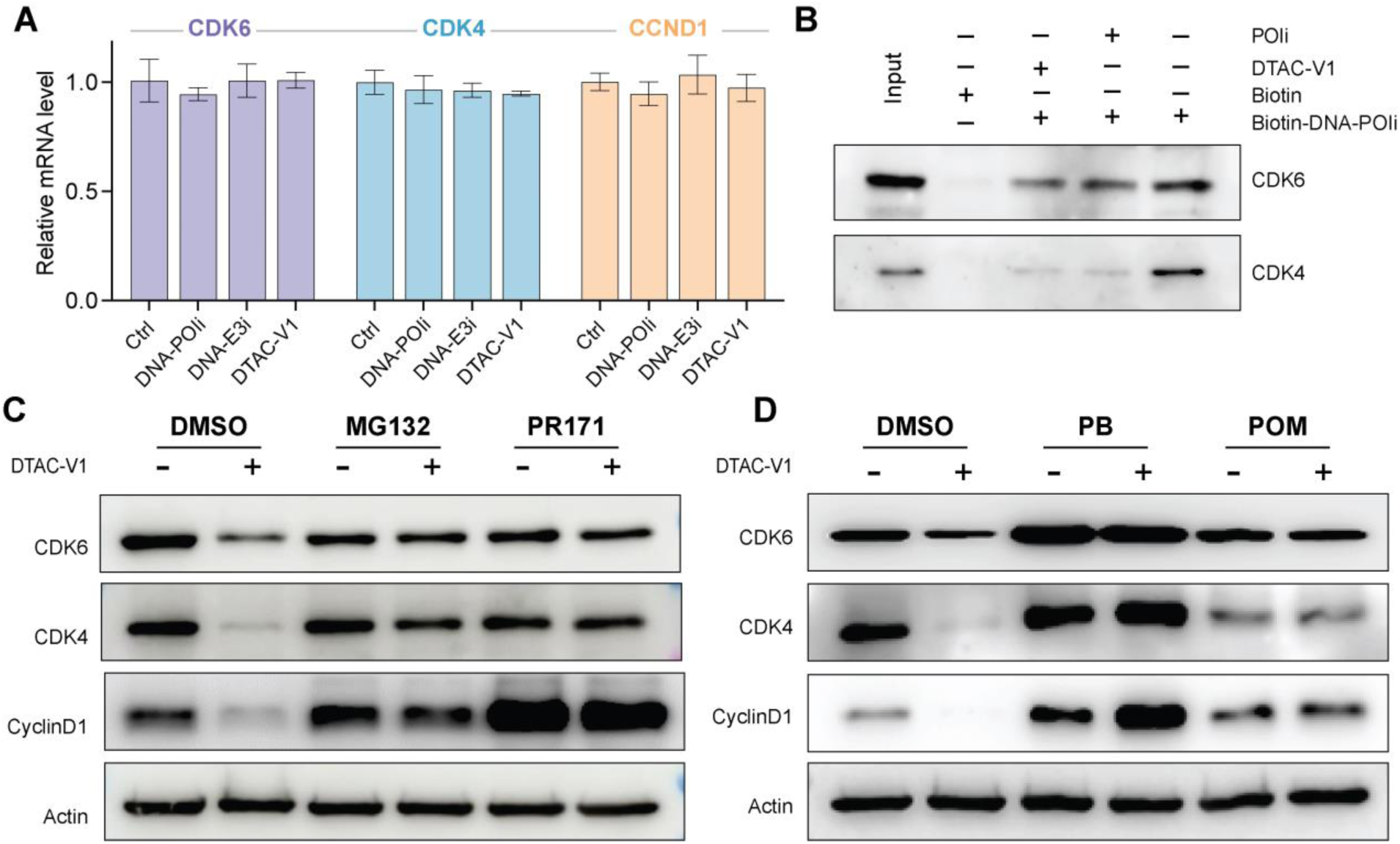
DTAC-V1 mediates Cyclin D1-CDK4/6 degradation via a UPS-dependent mechanism. **(A)** DTAC-V1 did not change the mRNA levels of CCND1(CyclinD1), CDK4, and CDK6 in RT-qPCR studies. U251 cells were treated with 100 nM of DNA-only control (Ctrl), DNA-POIi, DNA-E3i, or DTAC-V1 for 6 hours. mRNA levels were normalized to the control group. Data are representative of three independent biological replicates. **(B)** Streptavidin pulldown assay indicating that DTAC-V1 and POIi competes with Biotin-DNA-POIi for binding to CDK4 and CDK6. **(C)** U251 cells were pretreated with the proteasome inhibitors carfilzomib (PR-171, 2 μM) and MG132 (2 μM) prior to treatment with 100 nM DTAC-V1 for 8 hours, followed by immunoblot analysis of cell lysates. **(D)** Pre-incubation with pomalidomide (POM/E3i, 1 μM) or palbociclib (PB/POIi, 1 μM) inhibited DTAC-V1-induced degradation of the Cyclin D1-CDK4/6 complex in U251 cells. Results are representative of at least two independent experiments.

### DTAC-V1 induces proteasome-mediated degradation of CyclinD1-CDK4/6

To determine whether the DTAC-V1-induced downregulation of the Cyclin D1-CDK4/6 complex is mediated via the ubiquitin-proteasome system (UPS), we sought to elucidate the mechanism underlying this process. To begin, we conducted a streptavidin pulldown assay using a biotinylated DNA duplex conjugated with POIi (biotin-DNA-POIi). Both PB and DTAC-V1 were found to effectively compete with the biotin-DNA-POIi, thereby blocking the pulldown of CDK4 and CDK6 from the cell lysates. In contrast, the biotin-DNA-only control did not pull down CDK4 or CDK6. These results confirm the binding ability of DTAC-V1 to CDK4/CDK6 (**Figure 4B**). We next investigated whether DTAC-V1 promotes the formation of a ternary complex between cereblon (CRBN) and the Cyclin D1-CDK4/6 complex, thereby facilitating their proteasomal degradation. To test this hypothesis, cells were pretreated with either the POI ligand (PB) or the CRBN ligand (POM) prior to DTAC-V1 treatment. Pretreatment with PB or POM significantly attenuated the DTAC-V1-induced degradation of the Cyclin D1-CDK4/6 complex, suggesting that DTAC effectively recruits CDK4/6 and CRBN into a ternary complex required for ubiquitin-mediated degradation **(Figure 4D).** Finally, to confirm that DTAC-V1-mediated degradation proceeds via the UPS, we conducted rescue experiments using proteasome inhibitors MG132 and Carfilzomib (PR-171). Inhibition of the UPS effectively rescued the DTAC-V1-induced degradation of Cyclin D1-CDK4/6 **(Figure 4C)**, further confirming that its reduction is dependent on the proteasomal degradation pathway. Collectively, these findings demonstrate that DTAC-V1 induces the synchronous degradation of CDK4, CDK6, and Cyclin D1 through a UPS-dependent mechanism.

### DTAC-V1 induces selective degradation of the CyclinD1-CDK4/6 complex

Cyclin D1 is essential for both tumor initiation and maintenance and is frequently amplified and overexpressed in various cancer types^32^. Despite its critical role, Cyclin D1 has long been considered an ’undruggable’ target due to the absence of small-molecule binders, limiting its accessibility through conventional PROTAC strategies^33^. Existing CRBN-recruiting CDK4/6 PROTACs, such as BSJ-03-123, CP-10, and MS140, are unable to degrade Cyclin D1. Notably, DTAC-V1, developed using the same CDK4/6 inhibitor (PB) as these PROTACs, achieved robust degradation of Cyclin D1, CDK4, and CDK6 as shown in **Figure 5A**. To further evaluate the selectivity of DTAC-V1, we assessed its activity across various CDKs. Remarkably, DTAC-V1 selectively reduced the protein level of the CDK4/6-Cyclin D1 complex, while other CDKs, such as CDK1, CDK2, and CDK9, remained unaffected **(Figure 5B)**. To explore the selective degradation mechanism of DTAC-V1, we conducted label-free quantitative proteomic analysis on U251 cells. From this analysis, ∼6,300 proteins were identified. Principal component analysis (PCA) highlighted distinct protein expression profiles between the control and DTAC-V1 treated groups, which were clearly segregated (**Figure S13B**). Notably, in the treatment group, 21 proteins were upregulated, while 135 were downregulated (**Figure 5C**). Among the downregulated proteins, Cyclin D1, CDK4, and CDK6 were prominently reduced, whereas other cyclins and CDKs remained unaffected (**Figure 5C, Figure S13A**), corroborating previous WB data. Functional enrichment analysis of the differentially expressed proteins revealed their involvement in critical biological pathways, including cell division, cell cycle regulation, and mitosis (**Figure 5D**). Further protein-protein interaction analysis revealed the network of interactions among downregulated proteins and the Cyclin D1-CDK4/6 complex (**Figure S14**). These results underscore the potent and selective ability of DTAC-V1 to degrade the CDK4/6-Cyclin D1 complex.

**Figure 5.**
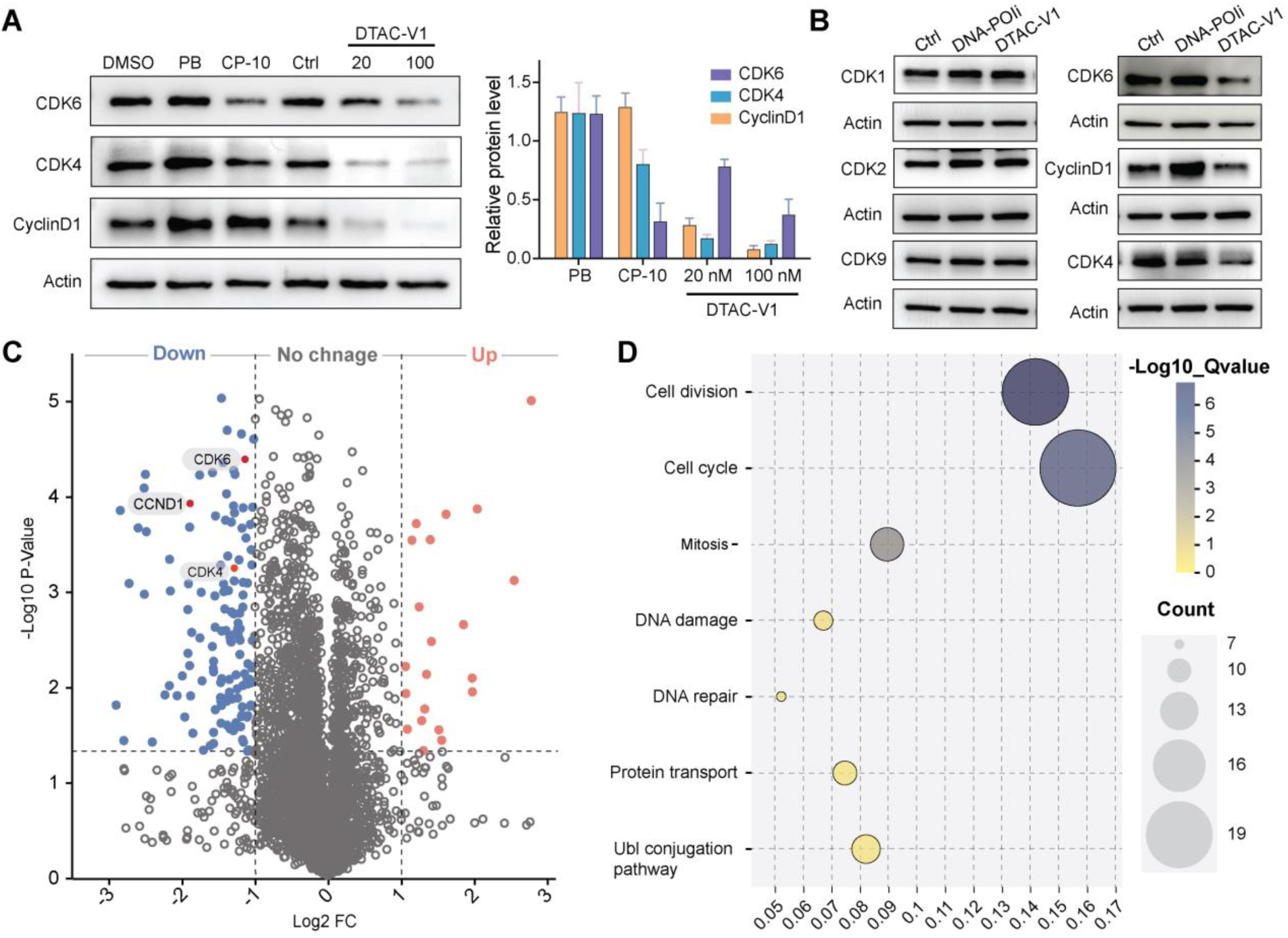
DTAC-V1 selectively degrades Cyclin D1 and CDK4/6. **(A)** *left:* western blot analysis of Cyclin D1, CDK4, and CDK6 levels in U-251 cells treated with PB (1 μM), CP-10 (1 μM), or DTAC-V1 (20 nM, 100 nM) for 16 h. Results are representative of at least two independent experiments. *Right:* quantification of relative Cyclin D1, CDK4, and CDK6 levels compared to the control group following the indicated treatments. Data are presented as mean ± SD from at least two independent experiments. **(B)** Western blot results showing DTAC-V1 selectively reduced Cyclin D1 and CDK4/6, but not CDK1, CDK2, or CDK9 in U-251 cells after 6 h of treatment at 100 nM. Results are representative of two biological repeats. **(C)** Volcano plot from label-free proteomic analysis of U-251 cells treated with 100 nM DTAC-V1 and DNA control for 16 h, showing global proteomic changes. Downregulated proteins (log2 FC < -1, p-value < 0.05) are highlighted in blue, upregulated proteins (log2 FC > 1, p-value < 0.05) in red. CDK4, CDK6, and Cyclin D1 (CCND1) are highlighted. **(E)** Gene ontology (GO) analysis of downregulated proteins. The color intensity reflects the P-value of each GO term, and the dot size corresponds to the level of gene enrichment.

### DTAC-V1 effectively suppresses the growth of U251 cells

Cyclin-CDKs dysregulation is a hallmark of cancer, with CyclinD-CDK4/6 serving as key drivers of the G1 to S phase transition via phosphorylation and inactivation of the retinoblastoma protein (RB)^34^. We next examined the impact of DTAC-V1-induced degradation of the CyclinD1-CDK4/6 complex in cancer cells. As DTAC-V1 concentration increased, RB phosphorylation levels progressively decreased (**Figure S15C**). Moreover, DTAC-V1 treatment significantly elevated the proportion of cells in the G1 phase compared to the control and the inhibitors only groups (DNA-E3i and DNA-POIi), suggesting that degrading the entire protein complex induces stronger cell cycle arrest than inhibiting CDK4/6 kinase activity alone (**Figure 6A, B**). DTAC-V1 potently inhibited U251 cell proliferation, whereas both DNA-POIi and DNA-E3i produced only minor effects (**Figure 6D**). In a dose-dependent cell viability assay, DTAC-V1 exhibited far more potent anti-cancer activity compared to D-POIi as shown in **Figure 6E**. Furthermore, DTAC-V1 treatment significantly suppressed colony formation, reducing it by ∼70% at 20 nM and over 95% at 100 nM, whereas DNA-POIi and DNA-E3i treatments showed only ∼30% inhibition as shown in **Figure 6C**. In summary, DTAC-V1 induced CyclinD1-CDK4/6 degradation displayed much stronger anti-proliferative and growth-suppressing effects than the parental inhibitors, offering potential therapeutic advantages in cancer treatment.

**Figure 6.**
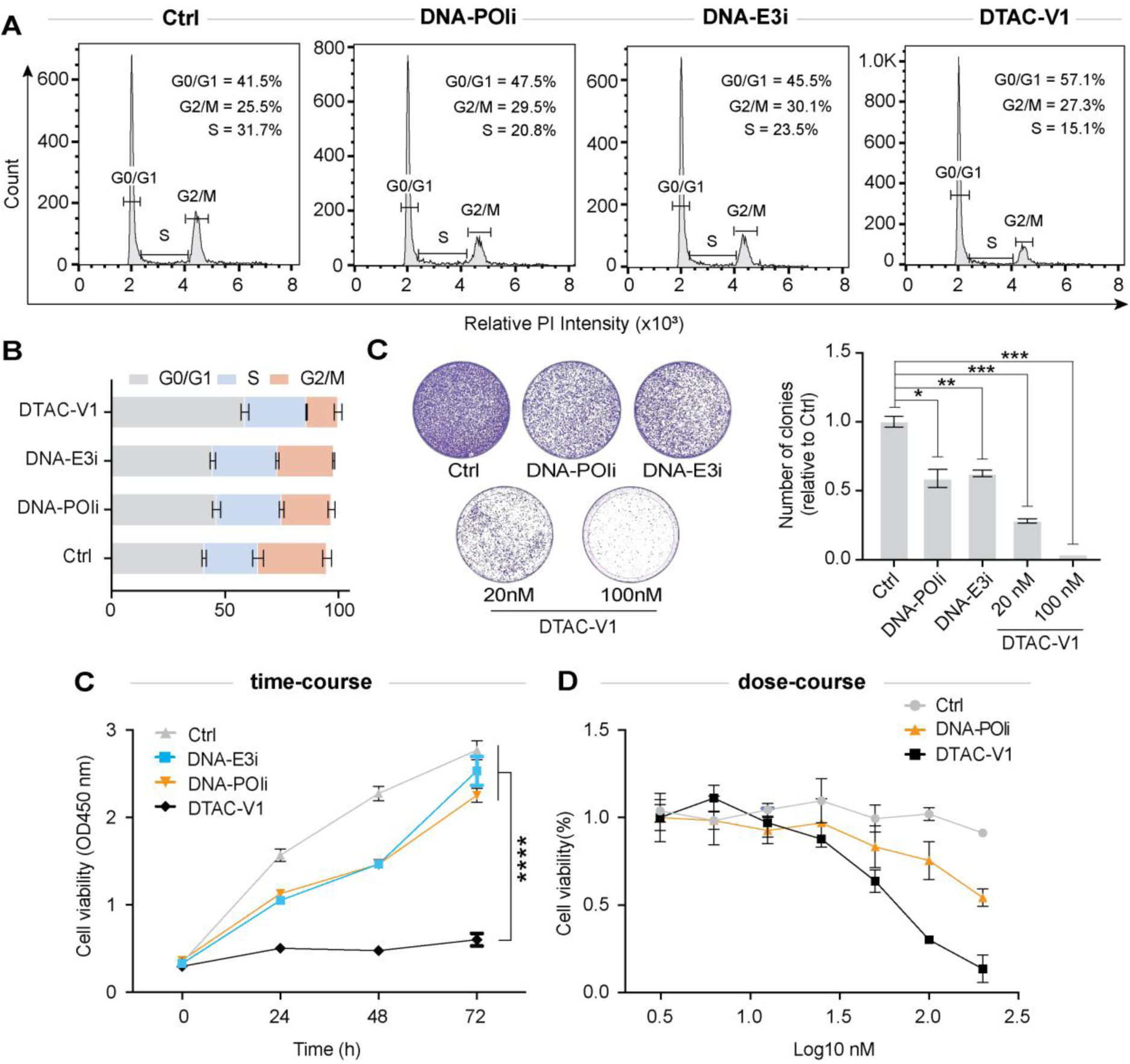
DTAC-V1 inhibits U-251 cell proliferation. **(A)** Flow cytometry-based cell cycle analysis of U-251 cells treated for 16 hours with either DNA control, DNA-POIi, DNA-E3i, or DTAC-V1. **(B)** Distribution of U-251 cells across G0/G1, S, and G2/M phases under the indicated treatment conditions. Data represent the mean ± s.d. of 2 to 4 biological replicates per condition. **(C)** *Left:* colony formation assay of U-251 cells treated with DNA control, DNA-POIi, DNA-E3i, or DTAC-V1. Representative images are shown. *Right:* quantification of U-251 colonies from (C) using ImageJ. Data are from two independent biological replicates. **(D)** Growth curves of U-251 cells treated with DNA control, DNA-POIi, DNA-E3i, or DTAC-V1 (100 nM for each treatment). **(E)** Cell viability y assay of U-251 cells after a 2-day treatment with DNA control, DNA-POIi, or DTAC-V1 at various concentrations. Error bars represent the mean ± s.d. from four biological replicates.

## Conclusion

In summary, we introduced a programmable DNA duplex-based PROTAC (DTAC) that demonstrated its potential to address significant challenges in the design and optimization of traditional PROTACs. By leveraging the programmability and structural control of the DNA scaffold, we achieved precise control of inhibitor spacing and orientation, resulting in distance- and orientation-dependent selectivity against the Cyclin D1-CDK4/6 complex. Notably, an optimal construct (DTAC-V1) led to its unprecedented and synchronous degradation without affecting other CDKs. This discovery emphasizes the specificity and efficiency of the design for targeting key regulatory proteins. DTAC-V1 further showed cell cycle arrest in the G1 phase, which significantly impaired cellular proliferation and effectively suppressed tumorigenic potential in cancer cells, highlighting its therapeutic promise in oncological applications.

Although this proof-of-concept work focused on the Cyclin D1-CDK4/6 complex, we aim to expand this technology as a general platform for both ex vivo and in vivo studies for discovery and functional analysis. This approach may be extended to other nucleic acid-based scaffolds such as clinically approved siRNA, or to synthetic nucleic acids such as threose nucleic acid (TNA), and L-DNA, which have been shown to enhance stability, pharmacodynamics, and to reduce immunogenicity^35–37^. Inspired by the programmability, adjustability, and modularity of DNA nanotechnology, future studies will focus on designing multifunctional scaffolds that target multiple proteins (either E3 ligases or target proteins) and could serve as versatile platforms for imaging, therapeutics, and cell targeting in concert^38,39^. The future holds great promise for this this technology, whereby it can be expanded for efficient screening of other proximity events such as post-translational protein modifications and lysosomal-mediated protein degradation^40,41^.

## Experimental procedures

### Resource availability

#### Lead contact

Further information and requests for resources should be directed to the lead contact, Hao Yan.

#### Materials availability

Reasonable requests for materials should be directed to the lead contact.

### Materials

Materials and detailed synthetic protocols for the synthesis of small-molecule inhibitors and ssDNA-inhibitor conjugates are provided in the supplementary file.

### Synthesis of DTAC versions

All DTAC constructs (V1, V2, V3, V4, V5, V6, V7, V8, and V9) were prepared by mixing complementary ssDNA-inhibitor conjugates (strand A and strand B) in equimolar ratio in 30 µL 1xPhosphate buffered saline (PBS, pH 7.4) buffer at 10-50 µM concentration. Thereafter reaction mixtures were annealed in a PCR thermocycler instrument using following protocol: Reaction mixtures were heated at 80°C for 3 min and then quickly cooled to 25°C.

### PAGE analysis of DTAC conjugates

Successfully synthesized DTAC library was analyzed using 12% native polyacrylamide gel electrophoresis (PAGE) as shown in **FigureS10**. To each lane was added 6 µL of a 2.5 µM of each sample, and the gel was electrophoresed at 90 V (constant voltage) for 1 h 30 min at 10°C, then stained with ethidium bromide (EtBr) and imaged with a Bio-Rad Molecular Imager GelDOC XR+ imaging system.

### Cell culture

Human glioma U251 cell line (Kindly provided by Dr.Yung.Chang) were cultured in DMEM (Gibco, Catalog No. 11965092) supplemented with 10% fetal bovine serum (BenchMark™,catalog No. 100-106-500 ) 1% antibiotics: penicillin (100 U/ml) and streptomycin (100 μg/mL;Gibco, catalog No. 15140122). All cell lines were grown at37 °C in a humidified 5%CO 2/ 95% air atmosphere. The number of living cells was calculated by Trypan Blue staining using a Countess 3 Automated Cell Counter (InvitroGen). Cell lines were regularly tested for mycoplasma contamination using Mycoplasma detection kit (InvivoGen, Catalog No. rep-mysnc-50).

### DTACs transfection

One day prior to DTACs transfection, cells were propagated into 6 cm cell culture dishes containing appropriate complete growth medium. Prior to transfection, complete medium was replaced with transfection medium (5%FBS, no Penstrep). DTACs transfection was performed using lipofectamine 3000 (Invitrogen, Catalog No. L3000001) reagent according to the protocols provided by the manufacture. All transfections were carried out in 6-cm dishes with 3 mL of media and concentrations of DTACs were calculated according to this volume (3 mL). Briefly, for 50 nM concentration, 5 µL from a 30 µM DTACs stock and 12 µL p3000 reagent was added to a tube containing 125 µL of Opti-MEM (Gibco, Catalog No. 51985034) and 8 µL of Lipofectamine 3000 reagent was added to a separate tube containing 125 µL of Opti-MEM. Two tubes were incubated for 5 minutes at room temperature and DTACs containing Opti-MEM was then slowly added to the second tube with lipo3000. The solution in the tube was mixed well by pipetting up and down several times. After incubating for 15-20 minutes at room temperature, 250 µL of DTACs-lipofectamine complex was added drop wise onto cells containing the transfection medium. Transfection medium was mixed well before transferring the plate into the incubator. After appropriate time, cells were either harvested or replaced with fresh medium and incubated for desired time point prior to harvesting.

### Immunoblotting and antibodies

Protein concentration in all the cell lysates were measured by BCA protein assay kit and equal amounts from each lysate were mixed with 4X loading dye and boiled for 5 minutes followed by 2 minutes centrifugation prior to loading into SDS-PAGE gel. Next proteins on the SDS-PAGE gel were transferred to a Nitrocellulose membrane by western blotting and the membrane was blocked with 5% milk in TBST (0.05%Tween 20) for 1 h. Primary antibodies were prepared in TBST with 2.5% milk and membranes were incubated overnight at 4°C. On the following day, membrane was washes for 15 minutes (Incubate for three times, 5 minutes each) and appropriate secondary antibodies (1:5000) were prepared in TBST and incubated with the membrane for 1h at room temperature (RT). Membrane was washed for 15 minutes with TBST (incubate for three times, 10 minutes each) prior to imaging.

The following primary antibodies were used in the study: β-Actin (Proteintech Group, Catalog No. 66009-1-Ig, 1:10000), β-Tubulin (Proteintech Group, Catalog No.66240-1-Ig, 1:10000), CDK6 (Cell Signaling Technology [CST], Catalog No.1331S,1:1000), CDK6(Abcam, Catalog No. AB124821, AB288368, 1:6000), CDK4(CST, Catalog No. 12790S, 1:1000; Abcam, Catalog No. AB108357, 1:3000), CDK9 (Abcam, Catalog No. AB76320, 1:5000), CDK1 (Proteintech Group, Catalog No. 10762-1-Ig, 1:4000), CDK2 (Proteintech Group, Catalog No. 10122-1-Ig, 1:5000),

### Immunofluorescence staining

U-251 cells were seeded into 10mm confocal dishes (Ibidi) and incubated overnight. Subsequently, cells were quickly washed thrice with ice-cold PBS and then fixed with 4% paraformaldehyde for 20min. After fixation, cells were washed with PBS and permeabilized with 0.1% Triton X-100 at room temperature for 10min. The samples were then incubated in a blocking solution for 1 h at room temperature, followed by incubation with primary antibody overnight at 4 °C. The primary antibody used was mouse CDK6 antibody (Abacm, 1:100). After washing thrice with PBST (PBS with 0.1% Tween 20), the samples were incubated with the secondary antibody in the dark for 1.5 h at room temperature. Cells were washed 3 times in PBST for 5 minutes each then incubated in the appropriate secondary antibodies supplemented CoraLite488-conjugated Goat Anti-Rabbit IgG(H+L) (Proteintech, 1:100) for 3 hrs. The nucleus was counterstained with DAPI (Hoechst 3342, 1ug/mL) for 20 mins followed by 3 times wash in PBST. Finally, the slides were Imaging by Nikon X confocal.

### Streptavidin Pulldown Assays

Cell lysates from U251 cells were incubated with biotin-DNA, biotin-DNA-POIi, DTAC-V1, or POIi (palbociclib) as indicated in **Figure 4B** at 4°C for 4 hours. 30 μL of streptavidin agarose beads were then added into the lysates and incubated at 4°C overnight. The beads were washed 6 times, boiled in 2 x SDS loading buffer, separated by 10% SDS-PAGE electrophoresis, and immunoblotted.

### Cell Proliferation and Colony Formation Assays

For cell proliferation assays, cells were seeded into 96-well plates and transfected with DNA only, DNA-POIi, DNA-E3i, or DTAC-V1 for the indicated times. Following transfection, CCK-8 kit substrate (Abcam, Catalog No. ab228554) was added to determine cell viability. For colony formation assays, U251 cells in a 24-well plate were transfected with DNA only, DNA-POIi, DNA-E3i, or DTAC-V1 for 12 hours. After transfection, the cells were further plated into a 6-well plate (1000/well). Two to three weeks later, the cells were fixed with 4% paraformaldehyde and stained with 0.4% crystal violet in 20% ethanol. The colonies were then imaged and quantified using ImageJ software.

### Cell cycle assay

U251 cells were seeded in 6-well plates and treated with 100 nM of either DNA-only control, DTAC-V1, DNA-POIi, or DMA-E3i for 14–16 hours. Following the treatment, the cells were harvested and fixed in ethanol. The fixed cells were then stained using the Cell Cycle Detection Kit (Abcam, Catalog No. ab139418) according to the manufacturer’s instructions. Flow cytometry data were analyzed using FlowJo software to assess the distribution of cells across the different phases of the cell cycle, including G1, S, and G2 phases.

### Quantitative Reverse Transcriptase Polymerase Chain Reaction (RT-qPCR)

U251 cells were seeded in 6-well plates at a density of 1.2 million cells per well. The following day, the cells were treated with 100 nM of DNA control, DNA-POIi, DNA-E3i, or DTAC-V1 for 6 hours. After the treatment, cells were harvested, and RNA was extracted using the Monarch Total RNA Miniprep Kit (New England BioLabs Inc., T2010S) according to the manufacturer’s instructions. Using 1 μg of total RNA, cDNA was synthesized with the SuperScript III First-Strand Synthesis System kit (Thermo Fisher Scientific, 18080051) following the manufacturer’s protocol. Gene expression was quantified using PowerUp SYBR Green Master Mix (Thermo Fisher Scientific, A25742) on the ViiA 7 Real-Time PCR system. The mRNA expression levels for each target gene were normalized to the internal control GAPDH and then calculated relative to the control group. All experiments were conducted in triplicate or quadruplicate and repeated three times. The human primer sequences used in the RT-qPCR analysis are listed in **Table S3**.

### LC−MS/MS Analysis

LC–MS/MS, solubilized proteins were quantified (Thermo Fisher EZQ Protein Quantitation Kit or the Pierce BCA), and 20ug total protein from each were processed using ASU Mass Spectrometry facility’s SOP. In brief, proteins were reduced with 50 mM dithiothreitol (Sigma-Aldrich) at 95°C for 10 min and alkylated for 30 min with 40 mM iodoacetamide (Pierce). Proteins were digested using 2.0 μg of MS-grade porcine trypsin (Pierce) and peptides were recovered using S-trap Micro Columns (Protifi) per manufacturer directions. Recovered peptides were dried via speed vac and resuspended in 30 μl of 0.1% formic acid. All data-dependent mass spectra were collected in positive mode using an Orbitrap Fusion Lumos mass spectrometer (Thermo Scientific) coupled with an UltiMate 3000 UHPLC (Thermo Scientific). Samples (1 uL) were direct-injected and fractionated using an Easy-Spray LC column (50 cm Å∼ 75 μm ID, PepMap C18, 2 μm particles, 100 Å pore size, Thermo Scientific). Electrospray potential was set to 1.6 kV and the ion transfer tube temperature to 300°C. The mass spectra were collected using the “Universal” method optimized for peptide analysis provided by Thermo Scientific. Full MS scans (375–1500 m/z range) were acquired in profile mode with the following settings: Orbitrap resolution 120,000 (at 200 m/z), cycle time 3 seconds and mass range “Normal;” RF lens at 30% and the AGC set to “Standard”; maximum ion accumulation set to “Auto;” monoisotopic peak determination (MIPS) at “peptide” and included charge states 2–7; dynamic exclusion at 60 s, mass tolerance 10 ppm, intensity threshold at 5.0e3; MS/MS spectra acquired in a centroid mode using quadrupole isolation at 1.6 (m/z); collision-induced fragmentation (CID) energy at 35%, activation time 10 ms. Spectra were acquired over a 180-min gradient generated using solution A, 0.1% formic acid/H2O, and solution B, 0.1% formic acid in 98% acetonitrile at a flow rate of 0.250 μl/min. Time course for gradient formation: 0–30 min at 2-15% B, 30-100 min at 15–28% B, 100–115 min at 28–35% B, 115–125 min at 35-45% B, 125-130 min at 45-98% B, 130-150 min at 45-98% B, 150-165 at 98-2% B, and 165-180 min at 2% B.

### Label-free quantification (LFQ) and Statistical Analysis

Raw spectra were loaded into Proteome Discover 2.4 (Thermo Scientific) and protein abundances were determined using Uniprot (www.uniprot.org) Homo sapiens database (Hsap UP000005640.fasta). Protein abundances were determined using raw files and were searched using the following parameters: Trypsin as an enzyme, maximum missed cleavage site 3, min/max peptide length 6/144, precursor ion (MS1) mass tolerance at 20 ppm, fragment mass tolerance at 0.5 Da, and a minimum of 1 peptide identified. Carbamidomethyl (C) was specified as fixed modification and dynamic modifications set to Acetyl and Met-loss at the N-terminus, and oxidation of Met. A concatenated target/decoy strategy and a false-discovery rate (FDR) set to 1.0% were calculated using Percolator. Accurate mass and retention time of detected ions (features) using the Minora Feature Detector algorithm were then used to determine the area-under-the-curve (AUC) of the selected ion chromatograms of the aligned features across all runs and the relative abundances calculated. Differential abundances between treatments were determined using protein abundance ratio t-tests (background based) as implemented in Proteome Discoverer 2.4^42^ and MaxQuant^43^.

### Data Processing and Bioinformatics Analysis

Differential gene expression analysis was conducted using the DESeq2 R package^44^. Genes with an adjusted p-value < 0.05 and |log2FoldChange| > 1 were considered significantly differentially expressed. Following this, Gene Ontology (GO) enrichment analysis was performed on the identified differentially expressed genes. Furthermore, all differentially expressed protein database accession or sequence were searched against the STRING database (v11.5, https://cn.string-db.org/) for protein–protein interactions.

### Statistical Analysis

Experimental data are expressed as the mean ± SD or SEM from three independent experiments, unless otherwise specified. Statistical significance between groups was assessed using a t-test in GraphPad Prism (version 8, GraphPad Software Inc., USA). A p-value of <0.05 was considered statistically significant. The levels of significance are indicated as follows: *P < 0.05, **P < 0.01, ***P < 0.001, ****P < 0.0001, and n.s. denotes no significant difference.

## Acknowledgements

We thank Chad Simmons for his valuable suggestions and assistance in editing the final version of manuscript. We would like to thank our undergraduate students Adam Doherty, and Vu Mai Thy Nguyen for their help in casting the in-house prepared western blot gels. We thank Ranjan Sasmal, Xinyi Tu, Liangxiao Chen, Hao Liu, and Lu Yu for helpful discussion. We would like to thank Dr. Timothy Karr, and Jessica Sandler from the Biosciences Mass Spectrometry/Proteomics Core at the Arizona State University for assisting in the proteomics study. We acknowledge the use of facilities within the Flow cytometry core, Advanced light microscopy core, and Magnetic resonance research center at the Arizona State University.

## Author contributions

R.Z. and A.P. contributed equally and will be putting their name first on the citation in their CVs. R.Z., A.P., Y.X., and H.Y. conceived the idea. A.P., R.Z., developed the experiments. R.Z., A.P., D.S. performed and interpreted all the experiments. A.P., R.Z. wrote the manuscript and made and edited the figures with input from H.Y. H.Y. oversaw the whole project, mentored, provided tremendous supports, and his insightful guidance.

