## Supplementary Text for "DNA-templated spatially controlled proteolysis targeting chimeras for CyclinD1-CDK4/6 complex protein degradation": YanLab_DTACs_Supplementary.pdf

**S1. Materials and supplies.**

Triethylammonium acetate buffer, methanol, 1-hydroxyadamantane, and dimethyl sulfoxide (DMSO) were purchased from Sigma Aldrich. The DBCO-sulfo-NHS linker was purchased from Glen Research (Fisher Scientific). Azido-PEG2-iodide (commercially listed as Azido-PEG3-iodide on the website) was purchased from Lumiprobe life science solution. Azido-PEG2-amine was purchased from Broadpharm.

**S2. *Smal molecule synthesis and characterizations***

*General Methods: All organic reactions were conducted in oven-dried glassware under argon or nitrogen atmosphere. Reagents were commercially available and used without further purification, while anhydrous solvents were procured as the highest grade from Sigma-Aldrich. The progress of reactions was monitored using thin layer chromatography (0.2 mm, silica gel 60 $F_{254}$). Small molecule inhibitors were purified using commercial silica gel column (40-60 mesh, 12g column size) on a flash column chromatography. Yields are presented as isolated yields of spectroscopically (NMR/HPLC) pure compounds. $^{1}H$ and $^{13}C$ NMR spectra were acquired using Bruker 400 MHz spectrometer. Chemical shifts are expressed in parts per million (ppm, δ) referenced to the residual $^{1}H$ resonance of the solvent (CDCl3, 7.26 ppm). $^{13}C$ spectra are referenced to the residual $^{13}C$ resonance of the solvent (CDCl3, 77.16 ppm). Splitting patterns are denoted as follows: s, singlet; br, broad; d, doublet; dd, doublet of doublets; t, triplet; q, quartet; m, multiplet.*

**4-((2-(2-(2-azidoethoxy)ethoxy)ethyl)amino)-2-(2,6-dioxopiperidin-3-yl)isoindoline-1,3-dione**
**(Azido-PEG2-pomalidomide; E3i)**

The **Azido-PEG2-pomalidomide** molecule was synthesized using previously published protocol with slight modification (**Figure S1**).[45] 4-Fluorothalidomide (0.100 g, 0.362 mmol, 1 equiv.), Azido-PEG2-amine (0.082 g, 0.471 mmol, 1.3 equiv.), and N,N-Diisopropylethylamine (189 µL, 1.086 mmol, 3 equiv.) were added into anhydrous dimethyl sulfoxide (1 mL) and reaction mixture was heated 100°C for overnight under argon atmosphere. Thereafter solvent was co-evaporated with ethanol several times. The crude mixture was purified on a flash chromatography, eluting with1:1 hexane:ethylacetate (50:50) to obtain yellow-orange solid powder as a product: yield (0.095 g, 0.220 mmol, 61.03%); $^1$H NMR (500 MHz, $CDCl_3$) δ 8.51 (s, 1H), 7.49 (dd, J = 8.5, 7.1 Hz, 1H), 7.10 (d, J = 7.1 Hz, 1H), 6.92 (d, J = 8.5 Hz, 1H), 6.50 (t, J = 5.6 Hz, 1H), 5.00 – 4.89 (m, 1H), 3.74 (t, J = 5.4 Hz, 2H), 3.69 (d, J = 4.1 Hz, 6H), 3.48 (q, J = 5.5 Hz, 2H), 3.38 (t, J = 5.0 Hz, 2H), 2.91 – 2.67 (m, 3H), 2.11 (ddd, J = 9.7, 7.2, 3.1 Hz, 1H) (**Figure S3**); QTOF-LC/MS (ESI$^+$): m/z calculated for $C_{19}H_{22}N_6O_6$: 430.16. $[M+Na]^+$; found 453.15.

**6-acetyl-2-((5-(4-(2-(2-(2-azidoethoxy)ethoxy)ethyl)piperazin-1-yl)pyridin-2-yl)amino)-8-cyclopentyl-5-methylpyrido[2,3-d]pyrimidin-7(8H)-one <u>(Azido-PEG2-palbociclib; POIi)</u>**

Palbociclib (0.100 g, 0.223 mmol, 1 equiv.), Azide-PEG2-iodide (0.083 g, 0.290 mmol, 1.3 equiv.), Potassium carbonate (0.077 g, 0.558 mmol, 2.5 equiv.), and catalytic amount of Tetraethylammonium bromide (0.015 g, 0.045 mmol, 0.2 equiv.) were added into anhydrous dimethylformamide (2 mL) and reaction mixture was heated at 90°C for 6 hours under argon atmosphere (**Figure S4**). After that reaction mixture was quenched with addition of distilled water (20 mL) and product was extracted with ethyl acetate (20 mL). Next, the organic phase was washed with brine, water and then dried over sodium sulfate for few minutes. Solvent was evaporated in reduced pressure and crude product was purified on a flash chromatography, eluting with 95:5 dichloromethane:methanol. Product was obtained as bright yellow solid: yield (0.07 g, 0.115 mmol, 52%) $^1$H NMR (500 MHz, $CDCl_3$) δ 8.86 (s, 1H), 8.62 (s, 1H), 8.15 (d, J = 9.1 Hz, 1H), 8.08 (d, J = 3.0 Hz, 1H), 7.33 (dd, J = 9.1, 3.0 Hz, 1H), 5.88 (q, J = 8.9 Hz, 1H), 3.68 (qd, J = 6.6, 3.6 Hz, 7H), 3.40 (t, J = 5.1 Hz, 2H), 3.22 (t, J = 5.0 Hz, 4H), 2.71 (dt, J = 15.8, 5.4 Hz, 5H), 2.55 (s, 3H), 2.38 (s, 5H), 2.17 – 2.00 (m, 2H), 1.96 – 1.83 (m, 2H), 1.75 – 1.58 (m, 2H), 1.26 (s, 1H) (**Figure S6**); QTOF-LC/MS (ESI$^+$): m/z calculated for $C_{30}H_{40}N_{10}O_4$: 604.32. $[M+H]^+$; found 605.33.

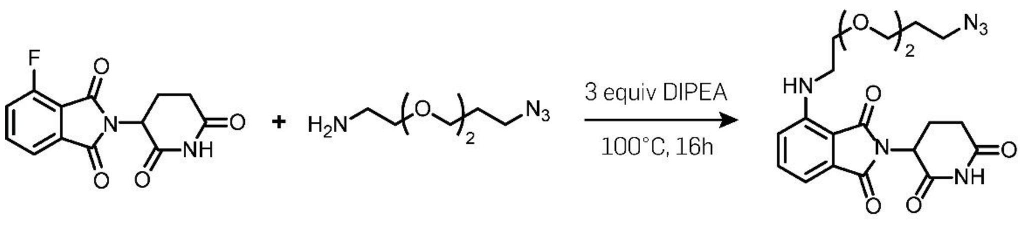


1

**Figure S1. Structure and chemical synthetic route for the Azido-PEG2-pomalidomide (E3i).** Details in experimental section.

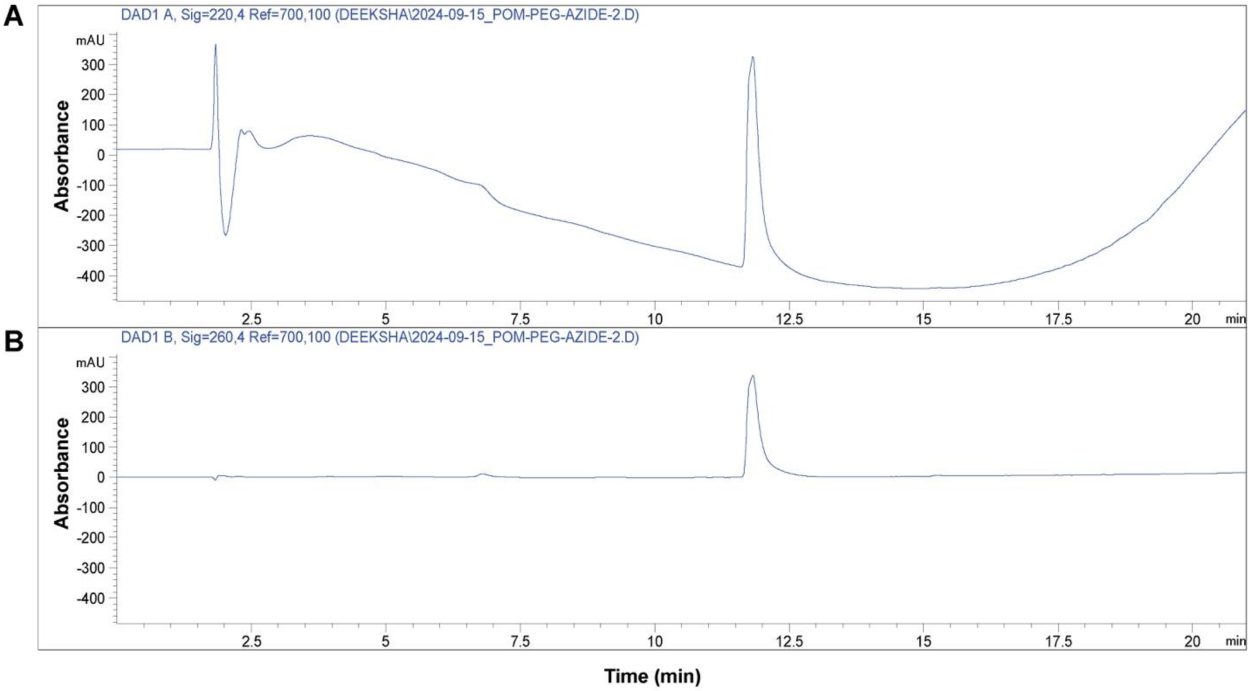


| Time (min) | Solvent A | Solvent B |
|---|---|---|
| 0 | 90 | 10 |
| 20 | 0 | 100 |
| 21 | 90 | 10 |

**Figure S2. Reversed phase HPLC trace of synthesized Azido-PEG2-pomalidomide:** (Solvent A: $H_2O$, Solvent B: ACN)

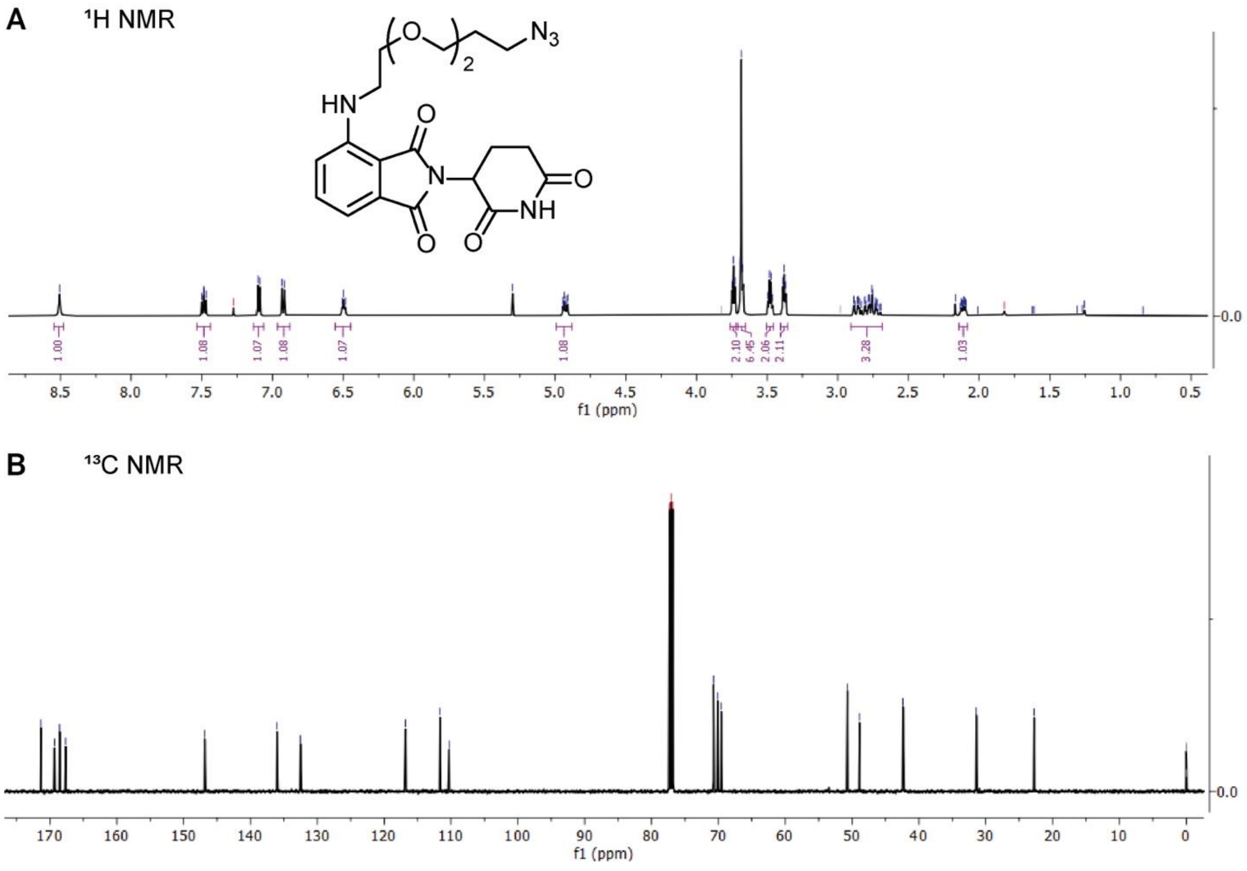


**Figure S3. NMR characterization of in-house synthesized Azido-PEG2-pomalidomide molecule (Solvent: $CDCl_3$).** A) 1H NMR. B) 13C NMR spectrum.

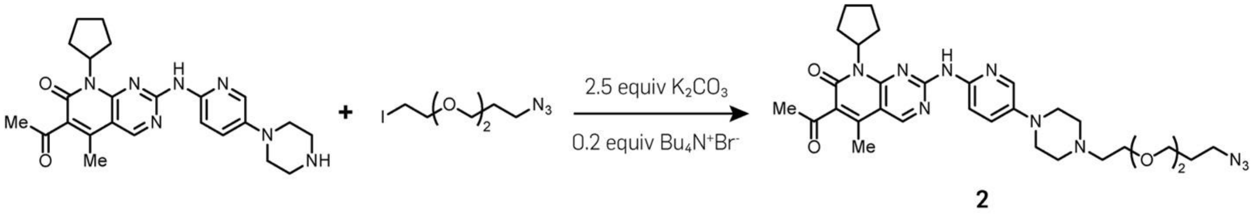


**Figure S4. Structure and chemical synthetic route for the Azido-PEG2-palbociclib (POIi).** Details in experimental section.

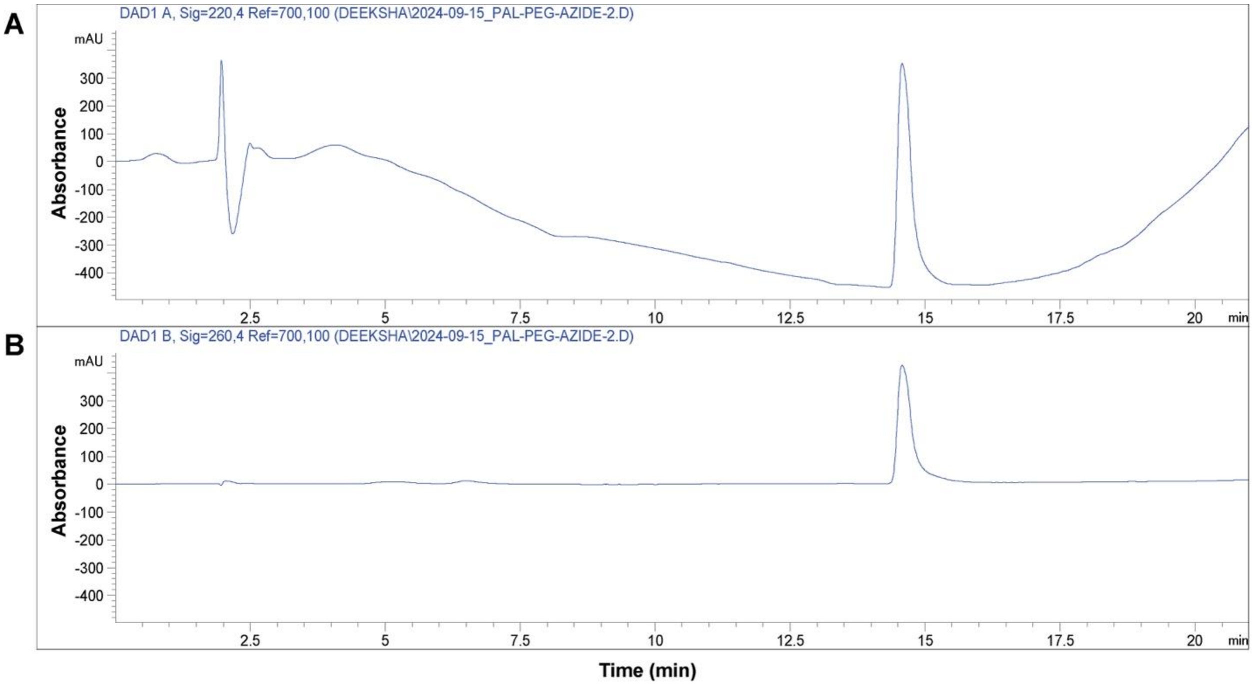


| Time (min) | Solvent A | Solvent B |
|---|---|---|
| 0 | 90 | 10 |
| 20 | 0 | 100 |
| 21 | 90 | 10 |

**Figure S5. Reversed phase HPLC trace of synthesized Azido-PEG2-Palbociclib:** (Solvent A: $H_2O$, Solvent B: ACN

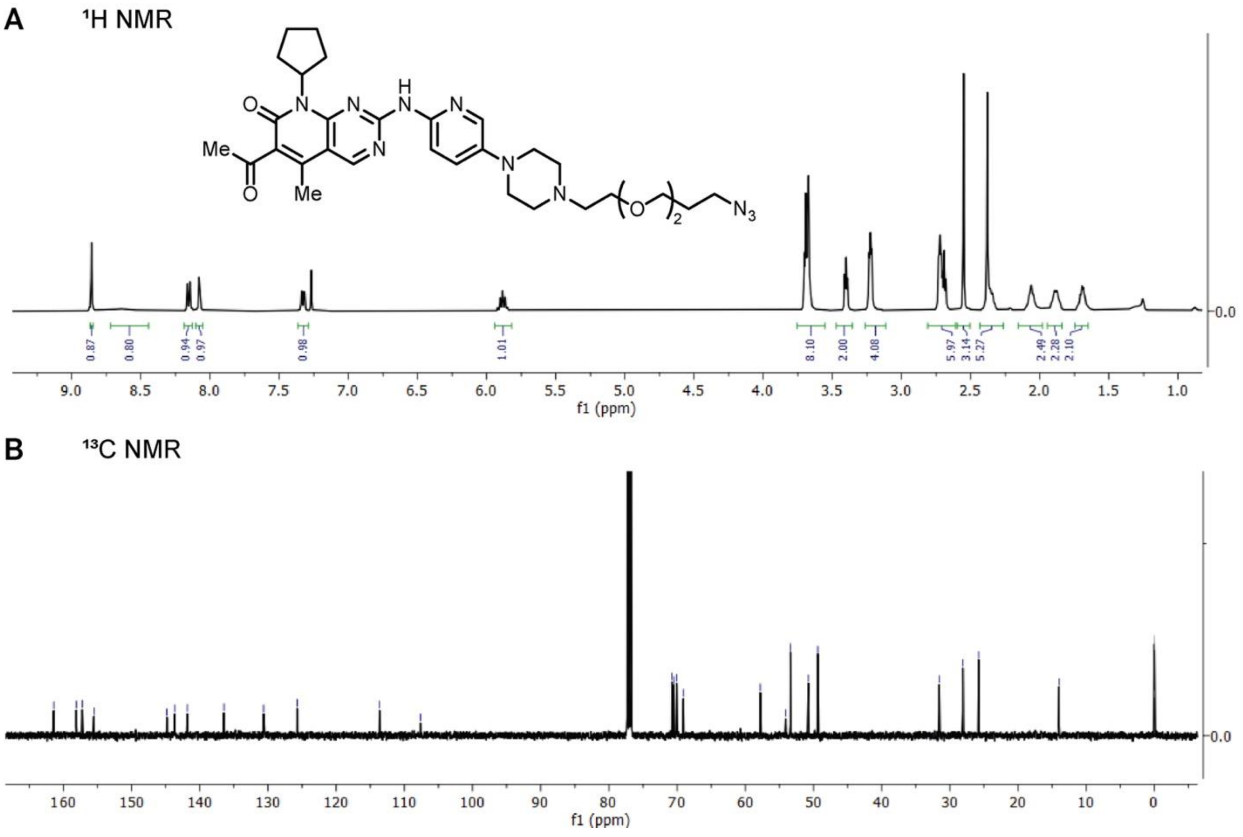


**Figure S6. NMR characterization of in-house synthesized Azido-PEG2-palbociclib molecule (Solvent: $CDCl_3$).** A) 1H NMR. B) 13C NMR spectrum.

***Amine modified oligonucleotides synthesis and characterization.***

The oligonucleotides were obtained via standard solid-phase oligonucleotide synthesis on a controlled pore glass (CPG, 1 μm). Standard DNA phosphoramidites, solid supports and additional reagents were purchased from Glen Research. The oligonucleotides were synthesized on an Applied Biosystems 3400 automated DNA/RNA synthesizer using a standard 1.0 μmole phosphoramidite cycle of acid-catalyzed detritylation activation and coupling, capping, and iodine oxidation. Stepwise coupling efficiencies and overall yields were determined by automated trityl cation conductivity monitoring. For phosphorothioate modification, 0.05M sulfurizing reagent II is prepared by dissolving in 40mL pyridine first, followed by 60mL acetonitrile to form a homogeneous solution. It is connected to the Auxiliary port on the DNA synthesizer and can be used similar to general iodine oxidation. For internal amine modification, amino-serinol phosphoramidite is dissolved in anhydrous acetonitrile to a concentration of 0.1M immediately prior to use. Cleavage of the oligonucleotides from the solid support and deprotection was achieved by exposure to 30% ammonia solution for 120 minutes at 55 °C on a heating-block. The cleavage solutions were diluted with water and ammonia was removed by washing with water using a 100kDa Amicon filter. The sequences of the oligonucleotides synthesized are tabulated (where "**i-NH$_2$**" denotes (**Table S1**) the internal amine and ***** represents phosphorothioate modifications) with the mass spectrometry data. The small molecule to be attached to each version is mentioned in brackets.

***dibenzocyclooctyne (DBCO) modified oligonucleotides synthesis and characterization.***

To conjugate DBCO to an oligonucleotide, an amine-modified DNA strand (Please see DNA sequences used) was treated with 6-fold excess of 200mM DBCO-sulfo-NHS-ester (dibenzocyclooctyne-sulfo-N-hydroxysuccinimidyl ester) in a 1XPBS buffer at pH ~8.5 (**Figure 10B**). The mixture was gently shaken for 3h at 37°C. The DNA-DBCO conjugates were purified by 3kDa amicon filter at 8000 rcf by repeated washing (5X) with distilled water to remove the excess small molecules and salts. Following filtration, the mixtures were purified using RP-HPLC and the conjugate peaks verified using ESI-MS.

***Synthesis, purification, and characterizations of small molecule-DNA conjugates.***

Conjugates were synthesized using strain promoted alkyne azide cycloaddition (SPAAC) chemistry (**Figure 10C**), like previous reports. Briefly, to DNA-DBCO conjugates (in 1xTAE-12.5mM $MgCl_2$, pH 7.5) was added 2 molar equivalents of 10 mM Azido-PEG3-pomalidomide (E3-i), or Azido-PEG3-palbociclib (POI-i) as a solution in DMSO. The reaction mixture was agitated and maintained at 37°C overnight. The DNA-drug conjugates were purified by 3kDa amicon filter at 8000 rcf by repeated washing (5X) with distilled water to remove the excess small molecules and salts. Following reaction, the mixture was purified using RP-HPLC and the conjugate peaks verified using ESI-MS.

***Purification by RP-HPLC***

Following reaction, ssDNA-inhibitor conjugates were purified using a C-18 column on an Agilent 1220 Infinity LC HPLC. The mixtures were purified using a linear gradient method, with Buffer-A (50 mM triethylammonium acetate (TEAA)) and Buffer-B (methanol). A linear gradient was run from 10% to 100% Buffer B over 60 minutes. Conjugates were monitored and collected based on 260 nm (for DNA) and 309 nm (for DBCO) absorbances. Collected fractions were lyophilized overnight until dry.

***Intact mass analysis of DTAC conjugates***

Collected peaks were tested for purity and identified by quadrupole time-of-flight liquid chromatography mass spectrometry (QTOF LC/MS) analysis in the negative mode. A 50 μM solution is prepared from the stock of ssDNA-inhibitor conjugates for the analysis using 0.1% ammonium hydroxide as the mobile phase. Identified peaks are deconvoluted in the expected mass range to get the intact mass of the conjugates (**Figure S8, S9**). The sequences of control ssDNA, and ssDNA-inhibitor conjugates are shown in **Table S1 and S2**.

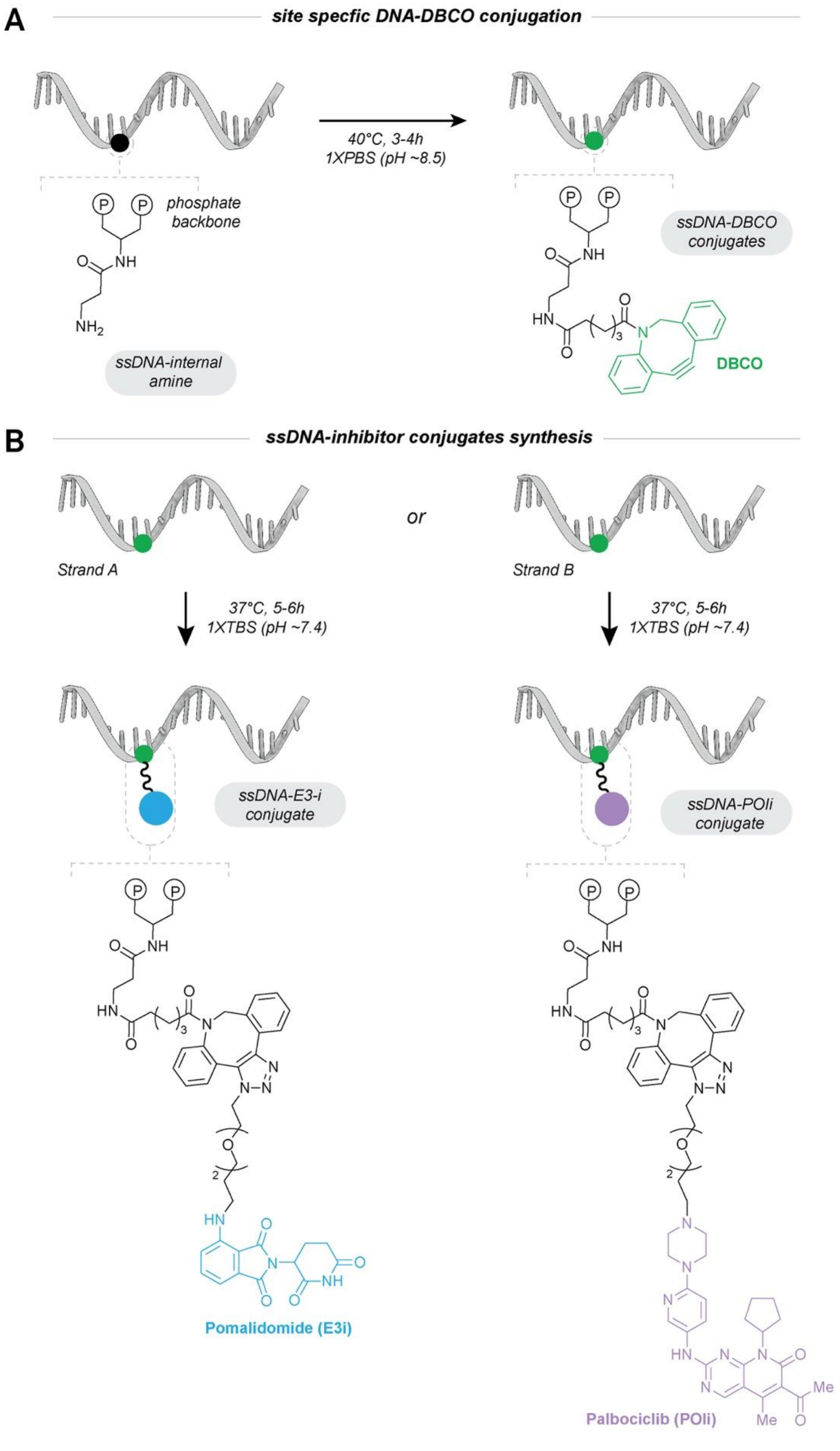


**Figure S7: Synthesis of DNA-drug conjugates.**

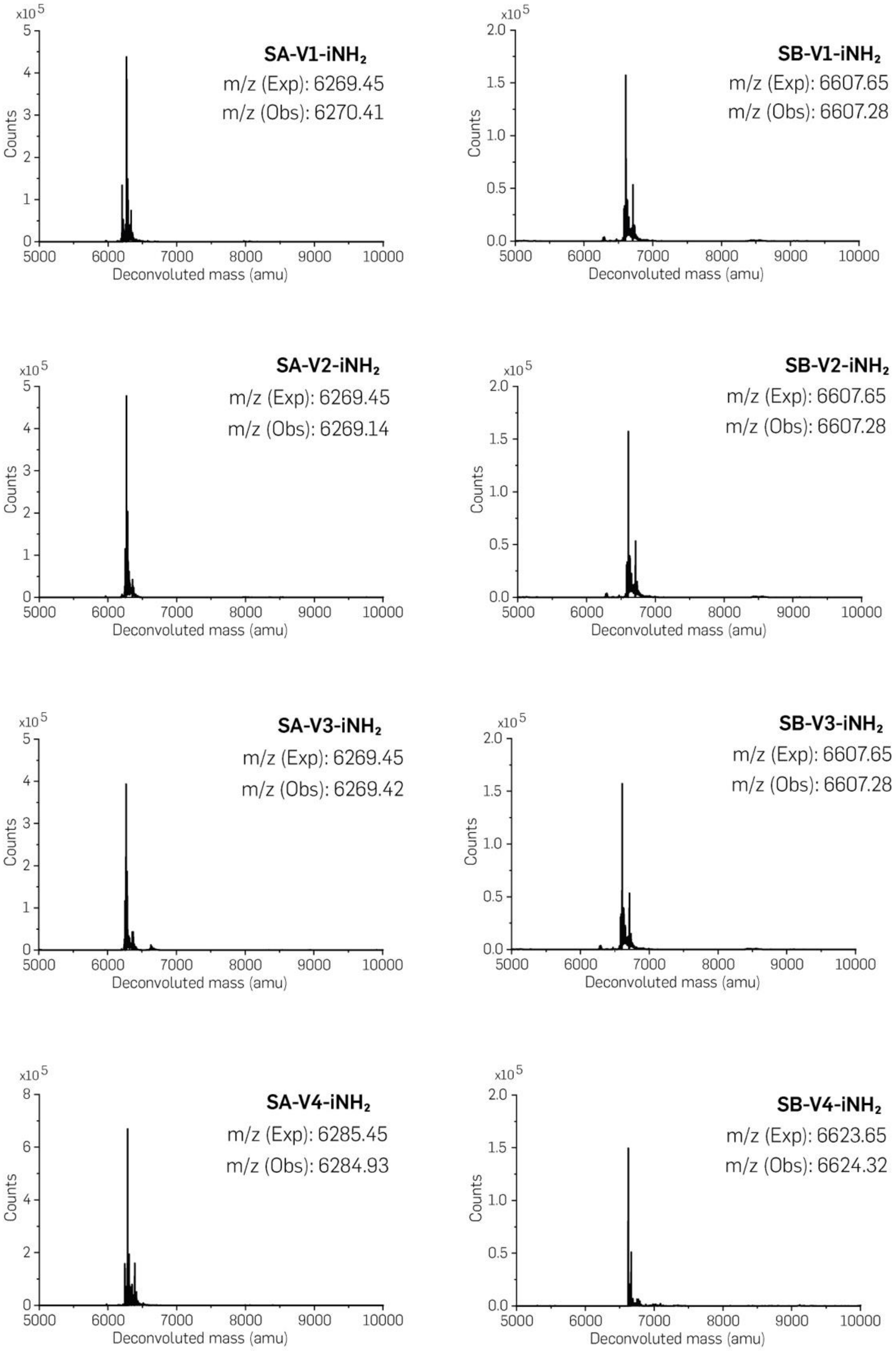


**Figure S8: Characterization of purified site specific amino modified ssDNA strand A&B.**

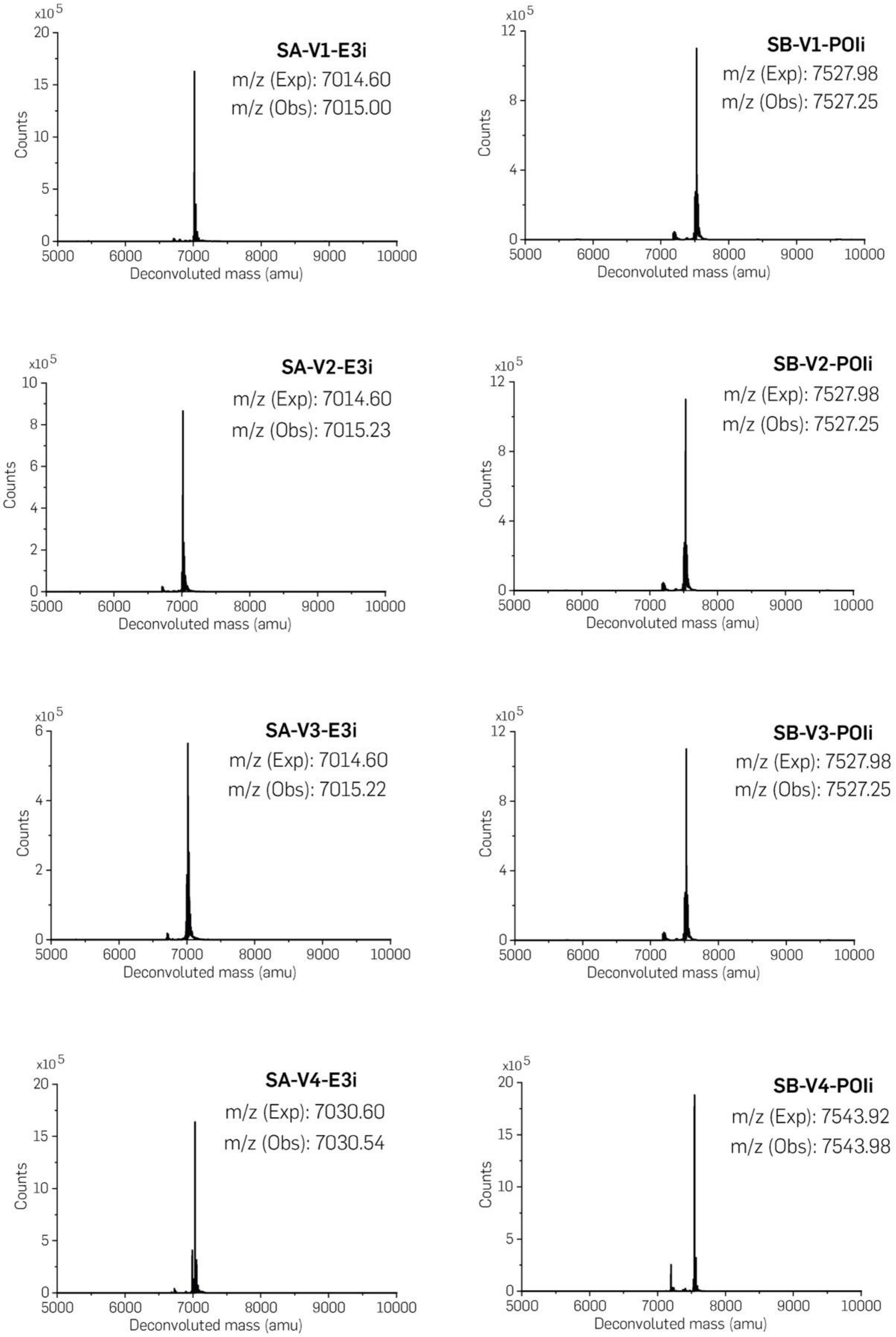


**Figure S9: Characterization of purified site specific E3i modified strand A and POIi modified strand B.**

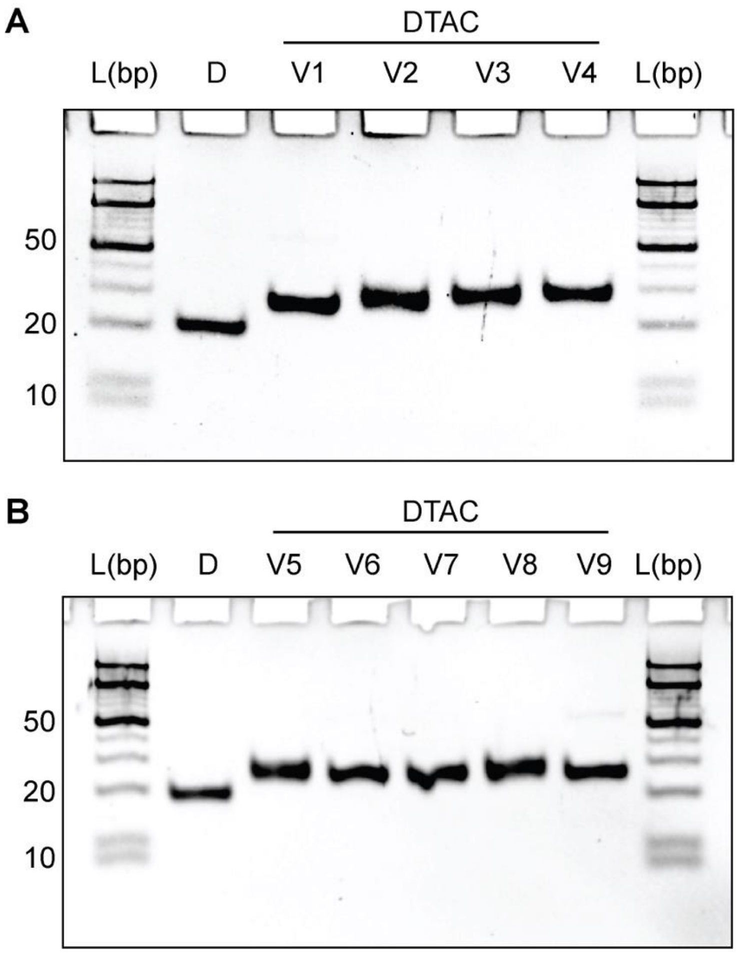


**Figure S10: Native PAGE (12%) gel electrophoresis of thermally annealed DTAC constructs. (A)** DTAC distance library**. (B)** DTAC orientational library.

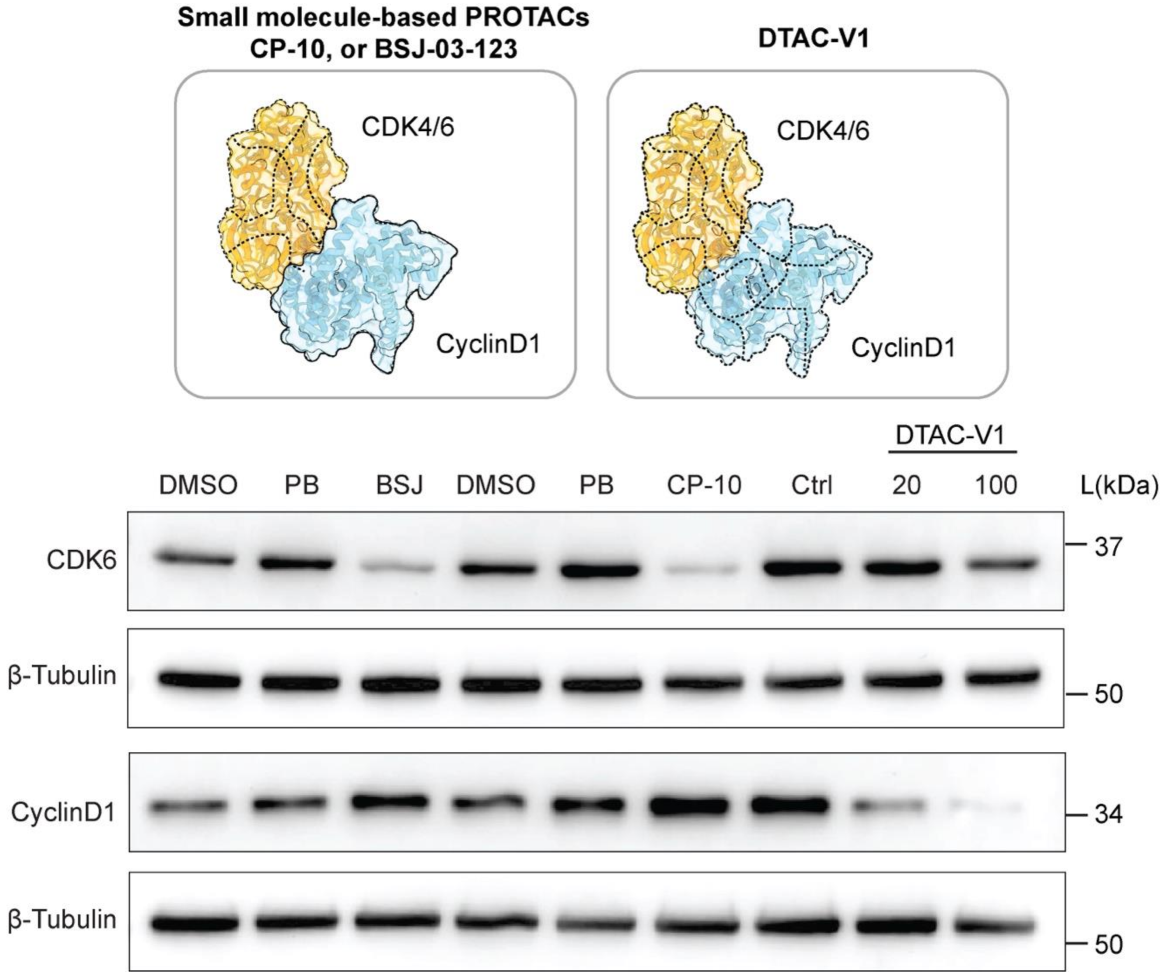


**Figure S11. Comparative effects of BSJ, CP-10, and DTAC-V1 on CDK6 and Cyclin D1 protein levels.** BSJ and CP-10 selectively induce degradation of CDK6 but not Cyclin D1, whereas DTAC-V1 depletes both CDK6 and Cyclin D1.

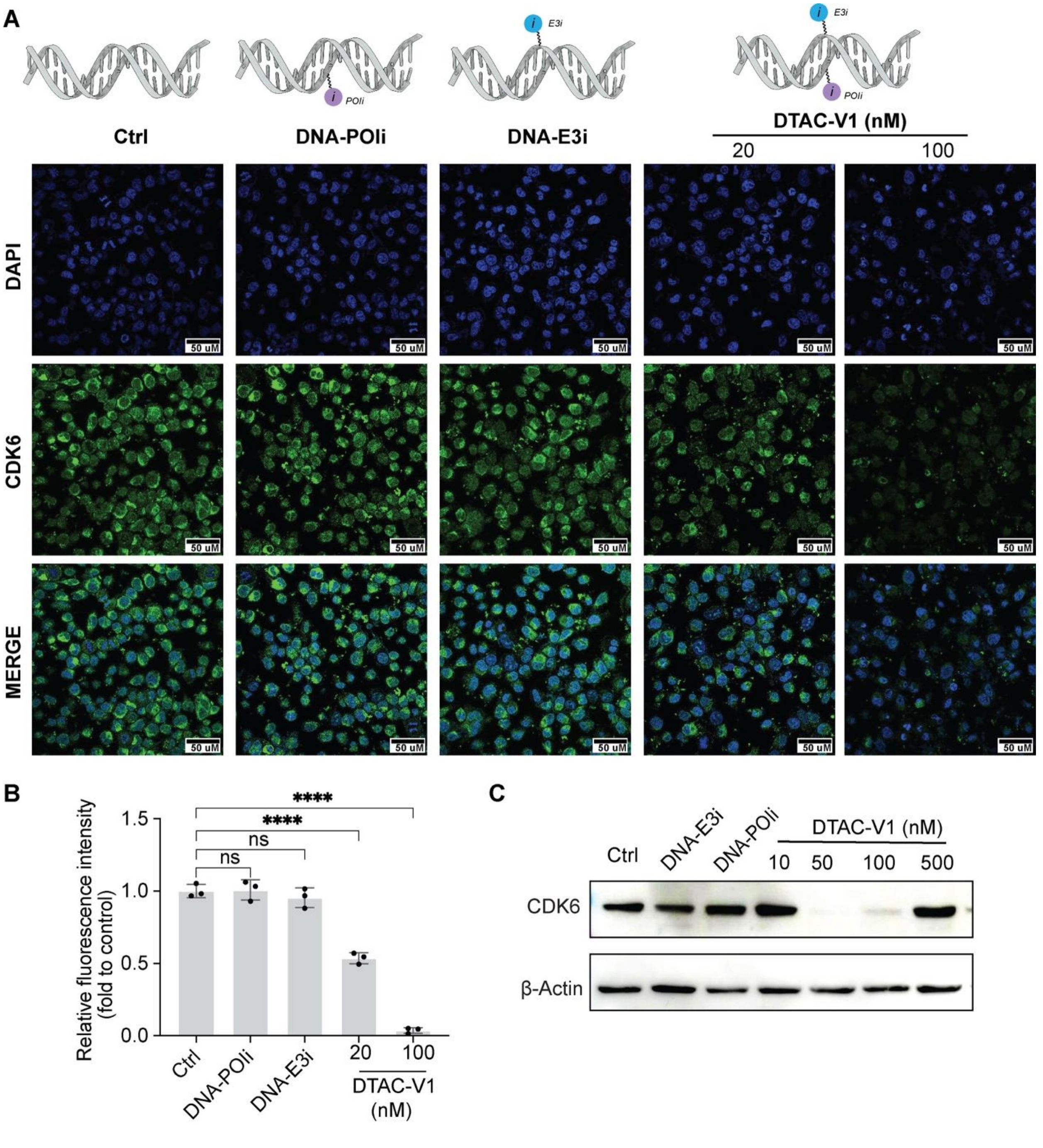


**Figure S12. Validation of DTAC-V1-induced CDK6 degradation via Western blot (WB) and immunofluorescence (IF). (A)** IF staining of CDK6 protein levels in U251 cells under the indicated treatments. Scale bar 50 μm. **(B)** Statistical analysis of relative fluorescence intensity of CDK6 in each treatment group. Statistical significance is denoted as ****$P < 0.0001$; n.s. denotes no significant difference. The horizontal lines above the bars indicate that the grouped bars share the same P-value. **(C)** WB analysis of CDK6 protein levels following treatment with DNA control, DNA-POIi (100 nM), DNA-E3i (100 nM), and DTAC-V1 at varying concentrations. A hook effect was observed at 500 nM DTAC-V1.

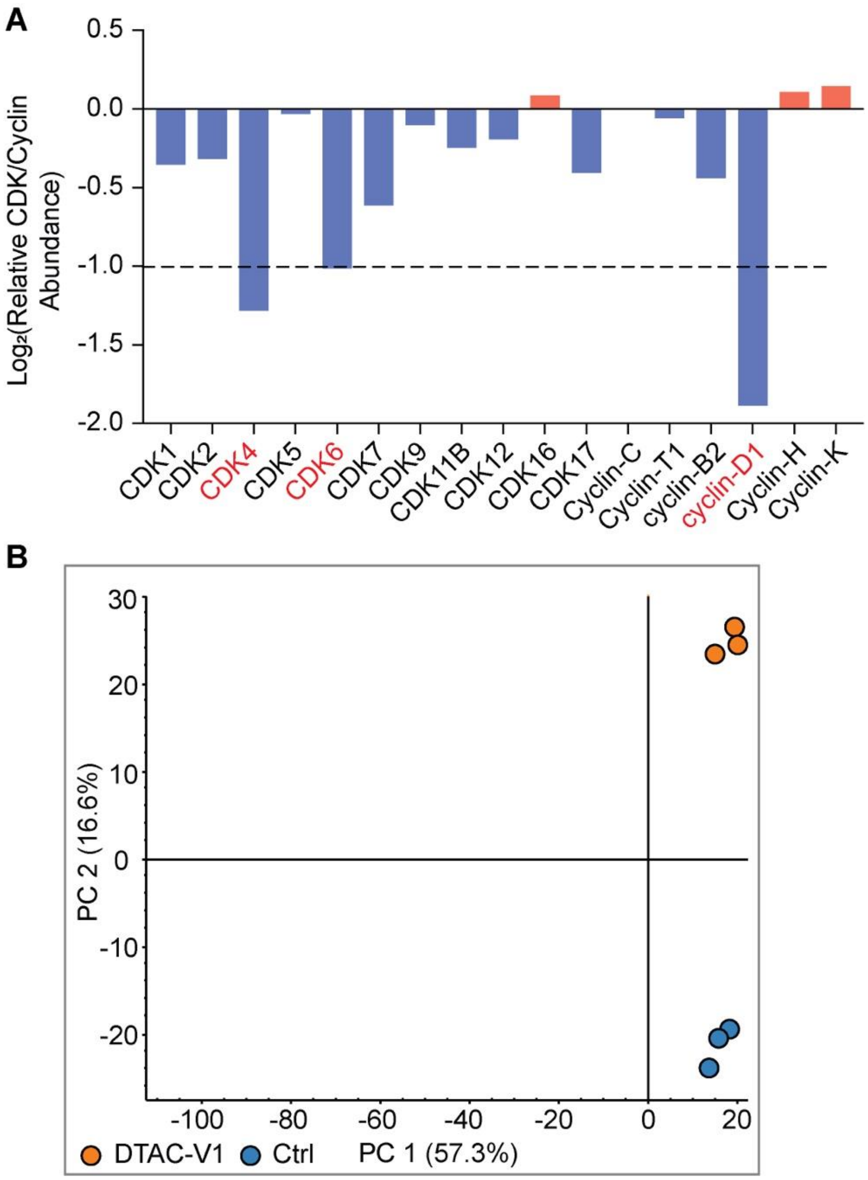


**Figure S13. Selective degradation of the CDK4/6-Cyclin D1 complex induced by DTAC-V1. (A)** Selectivity profile of DTAC-V1 on various CDKs and Cyclins as determined by proteomic analysis. **(B)** Principal component analysis (PCA) of U251 cells treated with DTAC-V1 compared to control groups. The PCA was performed on a dataset with replicates from three independent experiments (n=3), showing distinct clustering of the DTAC-V1 treatment group.

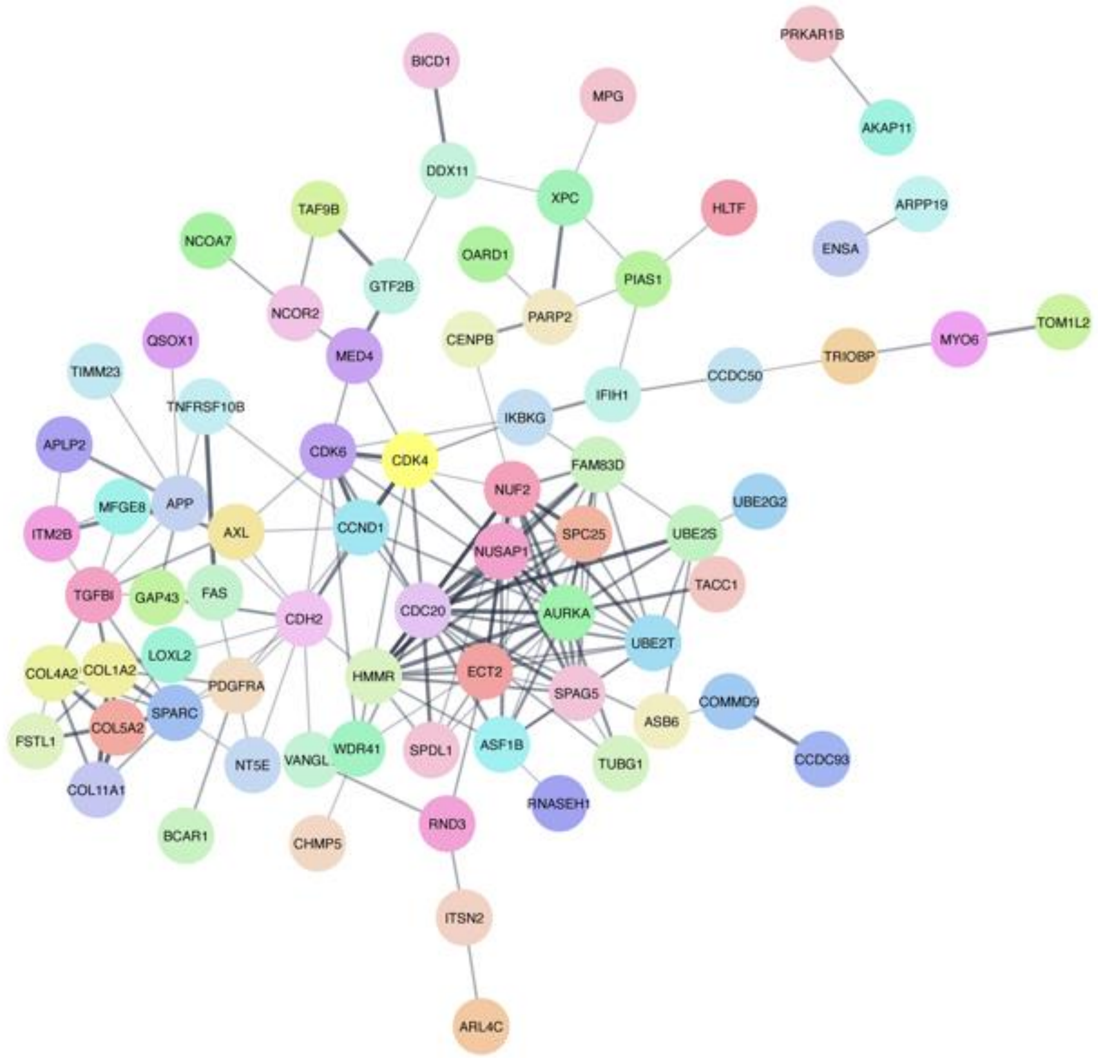


**Figure S14. Protein-Protein Interaction Analysis of Differentially Expressed Proteins in Proteomic Validation.** This figure illustrates the interaction network of proteins identified as differentially expressed in the proteomic analysis. Nodes represent individual proteins, and edges indicate interactions between them.

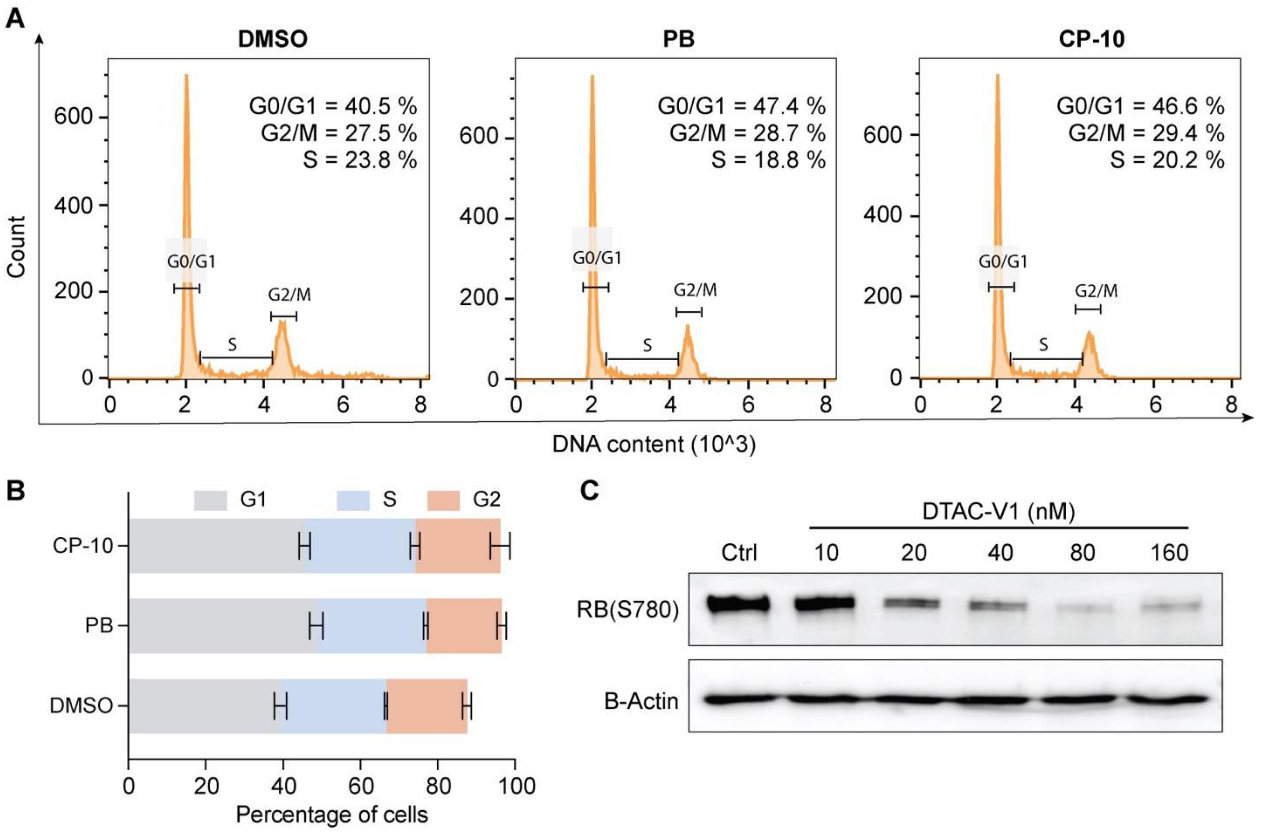


**Figure S15. Cell Cycle Distribution and RB Protein Phosphorylation Analysis.**

**(A)** Representative histogram showing cell cycle distribution after 20-hour treatment with the indicated concentrations (1 μM) of the specified compound. **(B)** Quantitative analysis of cell cycle distribution across treatments. Data are expressed as mean ± standard deviation from at least two independent biological replicates. **(C)** Western blot analysis demonstrating the effect of increasing DTAC-V1 concentrations on RB protein phosphorylation levels.

**Table S1**

X ≡ 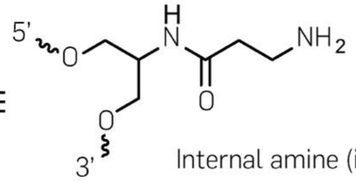

Internal amine ($iNH_2$)

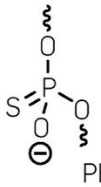

Phosphorothioate (*)

| Strand name | Sequence (5' to 3') | Expected mass (Da) | Observed mass (Da) |
|---|---|---|---|
| **SA-Ctrl** | **C*C*G*CAATCTCCTTCATC*G*C*C** | 5948.9 | 5948.1 |
| SA-V1-$iNH_2$ | C*C*G*CAATCTCXCTTCATC*G*C*C | 6269.45 | 6270.41 |
| SA-V2-$iNH_2$ | C*C*G*CAATCTCCTTCXATC*G*C*C | 6269.45 | 6269.14 |
| SA-V3-$iNH_2$ | C*C*G*CXAATCTCCTTCATC*G*C*C | 6269.45 | 6269.42 |
| SA-V4-$iNH_2$ | C*C*G*CAATCTCCTTCATC*X*G*C*C | 6285.45 | 6285.29 |
| **SB-Ctrl** | **G*G*C*GATGAAGGAGATTG*C*G*G** | 6287.1 | 6286.6 |
| SB-V1,V2,V3-$iNH_2$ | G*G*C*GATGAAGXGAGATTG*C*G*G | 6607.65 | 6607.28 |
| SB-V4-$iNH_2$ | G*G*C*GATGAAGGAGATTG*X*C*G*G | 6623.65 | 6624.32 |

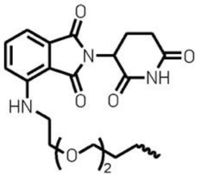

E3 ligase inhibitor (E3i)

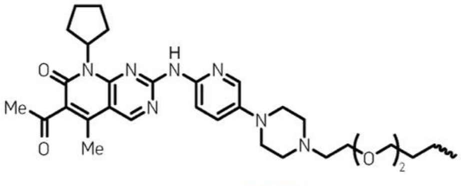

Protein inhibitor (POIi)

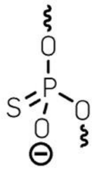

Phosphorothioate (*)

| | | | |
|---|---|---|---|
| SA-V1-E3i | C*C*G*CAATCTCE3iCTTCATC*G*C*C | 7014.60 | 7015.00 |
| SA-V2-E3i | C*C*G*CAATCTCCTTCE3iATC*G*C*C | 7014.60 | 7015.23 |
| SA-V3-E3i | C*C*G*CE3iAATCTCCTTCATC*G*C*C | 7014.60 | 7015.22 |
| SA-V4-E3i | C*C*G*CAATCTCCTTCATC*E3i*G*C*C | 7030.60 | 7030.54 |
| SB-V1,V2,V3-POIi | G*G*C*GATGAAGPOIiGAGATTG*C*G*G | 7527.98 | 7527.25 |
| SB-V4-POIi | G*G*C*GATGAAGGAGATTG*POIi*C*G*G | 7543.92 | 7543.98 |

**Table S2**

| Strand name | Sequence (5' to 3') | Expected mass (Da) | Observed mass (Da) |
|---|---|---|---|
| SA-V5-iNH$_2$ | C*C*G*CAATCTCCTTC**E3i**ATC*G*C*C | 6269.45 | 6269.14 |
| SA-V6-iNH$_2$ | C*C*G*CAATCTCCTT**E3i**CATC*G*C*C | 6269.45 | 6269.14 |
| SA-V7-iNH$_2$ | C*C*G*CAATCTCCT**E3i**TCATC*G*C*C | 6269.45 | 6269.14 |
| SA-V8-iNH$_2$ | C*C*G*CAATCTCC**E3i**TTCATC*G*C*C | 6269.45 | 6269.14 |
| SA-V9-iNH$_2$ | C*C*G*CAATCTC**E3i**CTTCATC*G*C*C | 6269.45 | 6270.41 |
| SB-V5,6,7,8,9-POli | G*G*C*GATGAAG**POli**GAGATTG*C*G*G | 7527.98 | 7527.25 |

**Table S3**

| Target gene | Forward primer | Reverse primer |
|---|---|---|
| CDK4 | ATGGCTACCTCTCGATATGAGC | CATTGGGGACTCTCACACTCT |
| CDK6 | TGCACAGTGTCACGAACAGA | ACCTCGGAGAAGCTGAAACA |
| CCND | GCTGCGAAGTGGAAACCATC | CCTCCTTCTGCACACATTTGAA |
| GAPDH | CTGGGCTACACTGAGCACC | AAGTGGTCGTTGAGGGCAATG |
